# Integrative transcriptomics reveals ectopic lipid homeostasis mechanisms in non-endocrine cells of the teleost pituitary

**DOI:** 10.1101/2021.06.11.448009

**Authors:** Eirill Ager-Wick, Gersende Maugars, Kristine von Krogh, Rasoul Nourizadeh-Lillabadi, Khadeeja Siddique, Romain Fontaine, Finn-Arne Weltzien, Christiaan Henkel

## Abstract

Directing both organismal homeostasis and physiological adaptation, the pituitary is a key endocrine gland in all vertebrates. It communicates the needs of the organism to different organs by secreting hormones into the bloodstream. Here, we have used the model fish medaka to investigate the developmental dynamics in the pituitary using a comprehensive RNA-seq time series. By linking developmental expression trends to single-cell RNA-seq profiles, we show how the transcriptional activities of cell types change during sexual maturation. One of the most prominent changes is the decline of the non-endocrine folliculo-stellate cell populations, and especially of rare cells expressing genes encoding secreted lipid transport proteins. As these genes are typically associated with the liver, this reveals the existence of unexpected connections between endocrine communication, lipid homeostasis, and sexual maturation.

## Introduction

All animals seek to grow, reproduce, and maintain homeostasis – but all at the right time. Several endocrine glands coordinate the present physiological needs of the organism and communicate these to different organs and tissues. The pituitary is a key endocrine gland common to all vertebrates, involved in the regulation of many important physiological processes, which it modulates by releasing several peptide hormones into the bloodstream. By integrating signals derived from the brain, as well as feedback signals from downstream peripheral organs, the pituitary holds the focal position in several endocrine regulatory systems, including the brain-pituitary-gonadal (BPG), brain-pituitary-thyroid (BPT) and brain-pituitary-adrenal/interrenal (BPA/BPI) axes^1^.

Over the lifespan of an animal, the pituitary gland exhibits a high degree of developmental plasticity, allowing the proliferation or reduction of specific cell types to meet changing demands^2^. Distinct cell types are responsible for the production and secretion of at least eight different peptide hormones^1,3^. Sexual maturation, for instance, is controlled by pituitary gonadotrope cells producing follicle-stimulating and luteinizing hormones (Fsh and Lh, respectively). These cells are themselves under the control of several brain-derived factors (e.g. the hypothalamic gonadotropin-releasing hormone, Gnrh) and subject to feedback through sex steroids produced by the gonads and other organs^4^. Understanding pituitary mechanisms is, therefore, a central goal of endocrinological research, as it provides a key handle on the experimental management of an animal’s physiology. In the case of teleost fish, this knowledge has important applications in aquaculture, ecology, and conservation.

One of the most important experimental windows on pituitary mechanisms is the transcriptome. Supported by high-throughput sequencing methods, it allows the sensitive inventory of a tissue’s investment in specific functions. As an early example of the potential of transcriptomics, the last major new peptide hormone discovered in the pituitary – the teleost-specific somatolactin – was identified based on cDNA only^5^. A current extension of transcriptomics is single-cell (sc) RNA-seq, in which transcriptome profiling of individual cells within a heterogeneous tissue allows the inference of cell type-specific mechanisms and developmental trajectories^6^.

To study the development of pituitary mechanisms over the course of sexual maturation, we here combine regular (bulk) RNA-seq and scRNA-seq on the pituitary gland of the model fish medaka, *Oryzias latipes*. By extracting expression trends from an RNA-seq time series of development, and mapping these to scRNA-seq profiles of mature fish, we show how cell types gain or lose prominence during sexual maturation (figure 1). This reveals two of the most striking changes to be the expected increase in maturation-related hormone production, and the unexpected expression and subsequent decrease of genes related to lipid homeostasis in non-endocrine folliculo-stellate cells. As these genes (including apolipoprotein A, the structural component of high-density lipoprotein particles) are typically only secreted by the liver, their unexpected and strong expression in the pituitary implies a major role in endocrine or paracrine communication.

**Figure 1.**
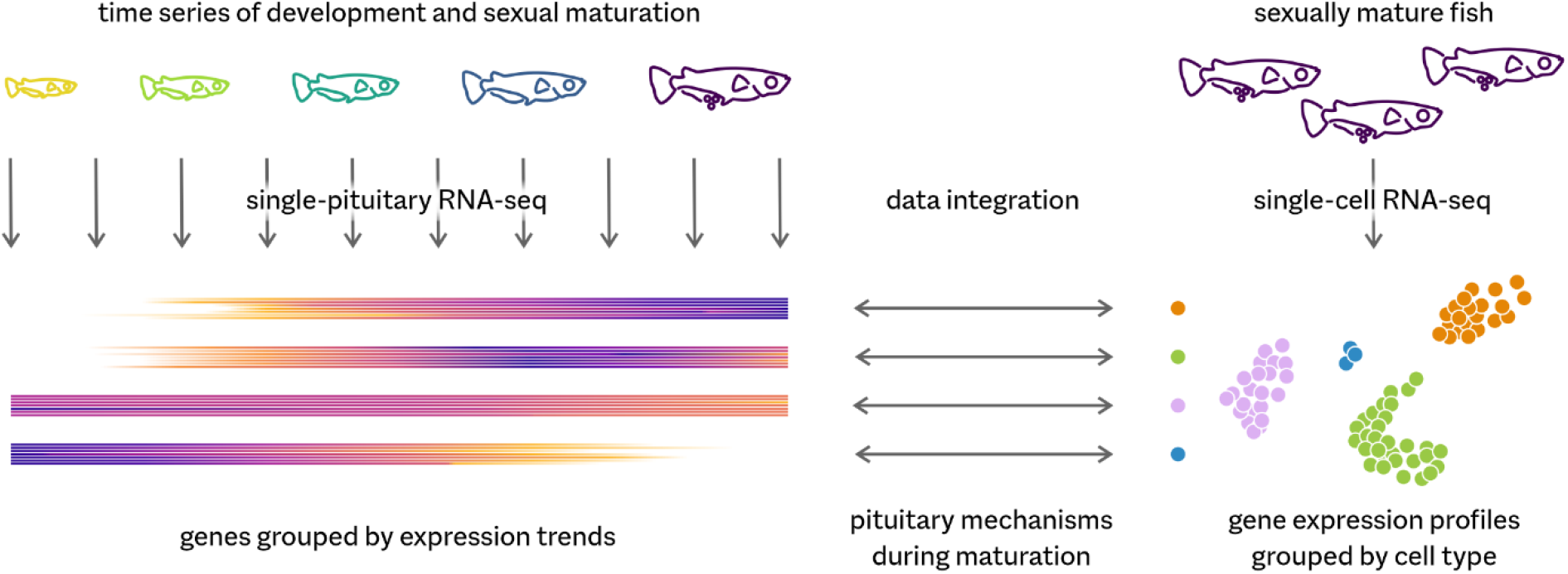
Study design. We infer gene expression trends during development and maturation from RNA-seq on pituitary glands of individual medaka fish. Using a previously described single-cell RNA-seq dataset on the pituitary of mature medaka, we can identify which cell types are associated with specific expression trends. In this simplified example, increase in gene expression in orange and green cells suggests they either proliferate or become more transcriptionally active during maturation; the expression trend associated with rare blue cells, on the other hand, implies they are more important during earlier developmental stages.

## Results

### A time series of pituitary development

In order to capture major changes in the pituitary during sexual maturation, we designed a time series experiment in which we bred medaka siblings (offspring from three couples) under controlled conditions. We sampled the pituitaries and ovaries of female fish after 31–178 days post-fertilization (dpf). The youngest fish were juveniles, and the smallest from which the pituitary gland can be reliably dissected; fish were fully sexually mature well before the final age. In addition, we sampled three older fish (249–277 dpf) from the same facility and the three parent fish (375 dpf). For all fish, we used the gonads for a histological assessment of maturation, and individual pituitary glands for profiling gene expression using RNA-seq (9.5–16.2 million reads per sample).

We subsequently selected a group of 54 fish for which growth correlates well with age (figure 2a, table 1, supplementary table 1), and focused on genes that show medium to high expression in the pituitary. As a selection criterion, we took the appearance in any sample in the top 1000 by gene expression level. Over all samples, this results in a set of 2686 medium to high expression genes (supplementary table 2), which suggests samples differ considerably by expression profile.

**Table 1.**
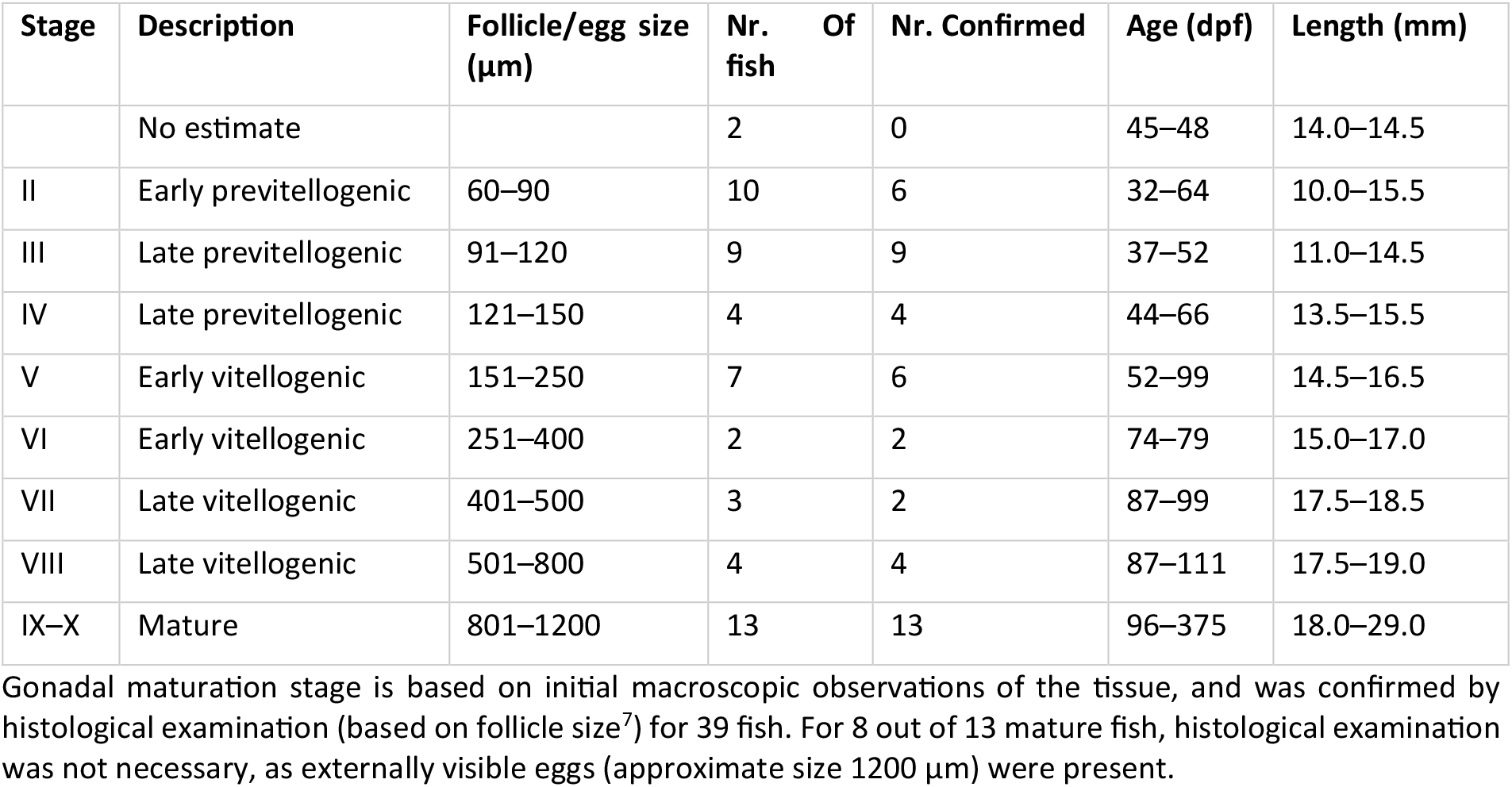
Sampled fish per gonadal maturation stage.

**Figure 2.**
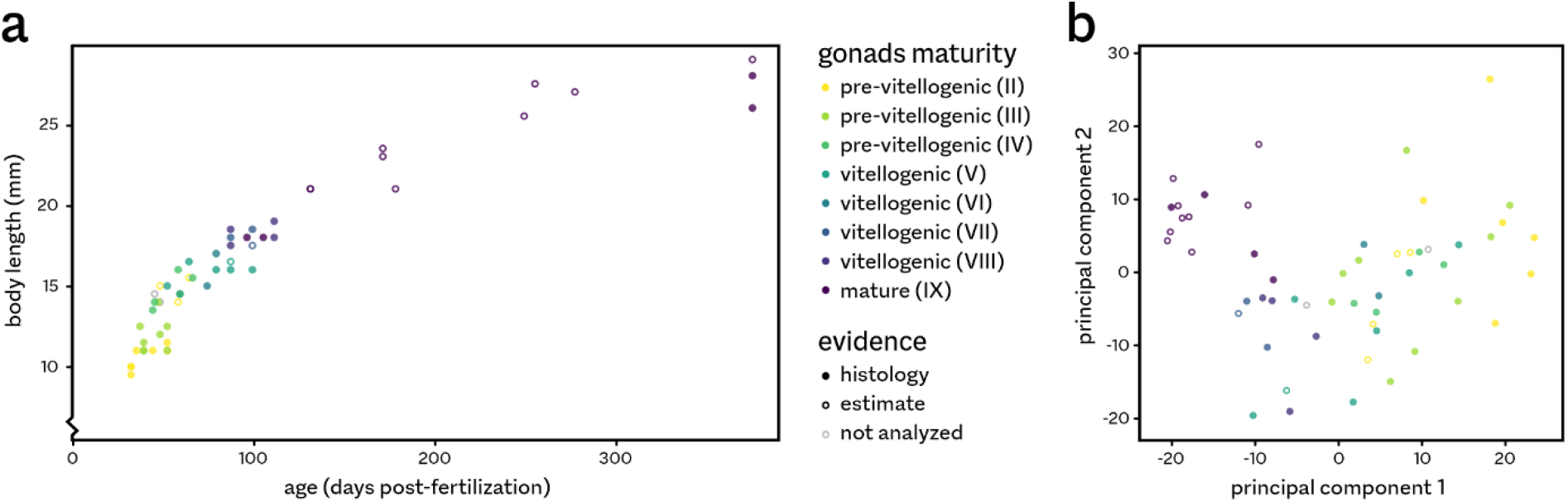
Growth and maturation in female medaka. **a** The 54 fish used in this study range from young, small and sexually immature (pre-vitellogenic gonads) to old, large and sexually fully mature. Gonad maturation status was assessed using histology for most samples; for a few fish, we only obtained macroscopic estimates. We did not perform histological staging for the gonads of most of the mature, spawning fish. **b** Principal component analysis of the pituitary transcriptomes shows a strong correlation of the first principal component (PC1) with maturation. PC1 explains 21.9% of between-sample variance, PC2 11.9%.

To investigate whether these differences are linked to pituitary development over time, we performed a principal component analysis (figure 2b), which identifies the most important sources of expression variation among the samples. The main trend (principal component 1, explaining 21.9% of variance) indeed correlates well with sexual maturation. We therefore decided to use this first principal component (PC1) as a measure of developmental ‘pseudotime’, which can be used to unambiguously rank the pituitary transcriptomes (supplementary figure 1). Of the 2686 medium-high expression genes, 52.7% are differentially expressed (5% false discovery rate) along this pseudotime axis, compared to 0.8% for the second principal component, and 36.4% and 37.3% with age and length, respectively. When grouped by developmental stage, 47.6% of genes are differentially expressed between pre-vitellogenic (combined stages II–IV) and mature (IX) fish, 14.6% between pre-vitellogenic and vitellogenic (V–VIII), and 28.0% between vitellogenic and mature.

### Divergent gene expression trends during sexual maturation

The genes encoding the pituitary peptide hormones are amongst the most highly expressed throughout development (supplementary figure 2). Both *fshb* and *lhb*, encoding the beta subunits of follicle-stimulating hormone and luteinizing hormone, respectively, show a significant increase in expression over time, as does the common alpha subunit *cga. pomca* (encoding the hormone precursor pro-opiomelanocortin) and *prl* (prolactin) are consistently the most highly expressed genes in the entire pituitary. The levels of *gh* (growth hormone) and *tshba* (thyroid-stimulating hormone beta subunit) increase slightly in mature animals, and *smtla* (somatolactin) expression decreases during development.

Figure 3 shows how these genes share several similar expression patterns over the course of development, which can be an indication of common regulatory mechanisms or functions. We used weighted gene co-expression network analysis^8^ (WGCNA) to categorize all 2686 medium-high expression genes into 24 expression modules, each representing a distinct expression trend (figure 4 and supplementary figure 3).

**Figure 3.**
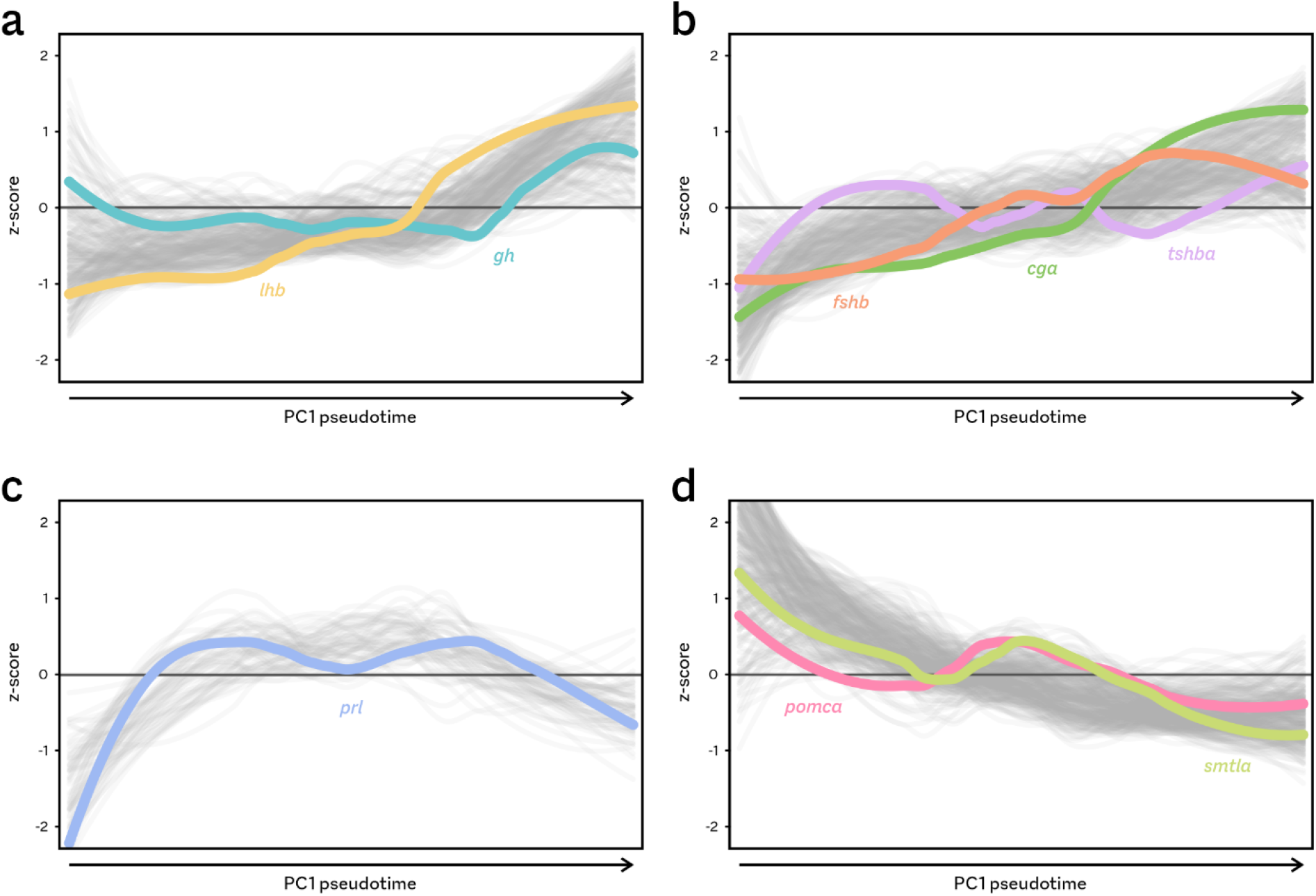
Hormone gene expression trends. Expression changes are shown as z-scores (standard deviations around the mean expression level) over development. Coloured lines indicate the expression trends (loess local regression) for hormone-encoding genes, grey lines for other genes sharing that pattern. The four panels represent distinct trends identified by WGCNA analysis, corresponding to modules in figure 4 and supplementary figure 3: **a** module 4, **b** module 6, **c** module 14, and **d** module 18.

**Figure 4.**
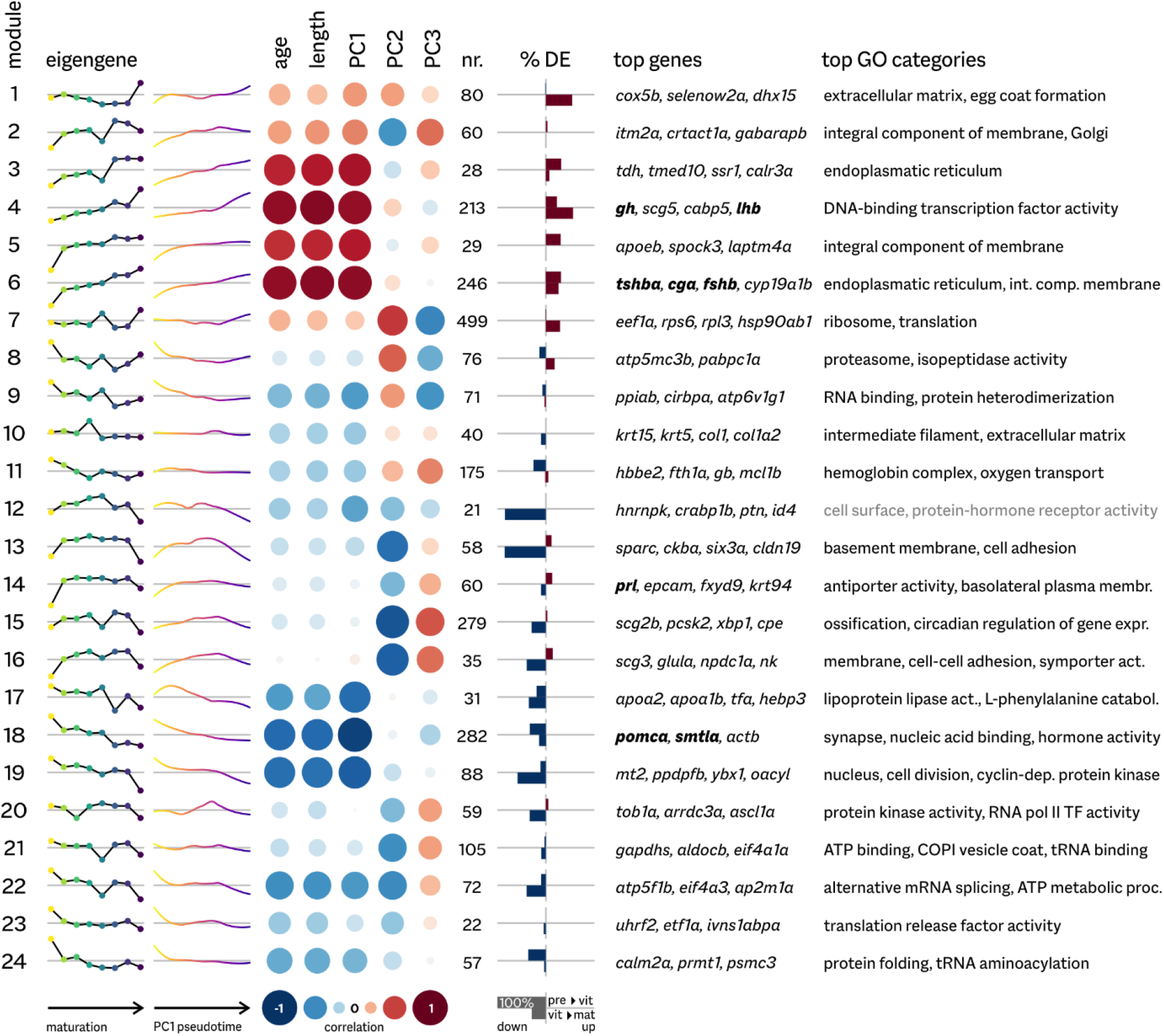
Expression modules. For each module, the typical expression pattern (eigengene) is shown both across gonad maturation stages and along principal component 1 (see figure 1b). These trends are based on variation around the mean expression over all samples (horizontal grey lines). Also shown are Spearman rank correlation with fish age and length, and with additional principal components; the number of genes in the module; the fractions of genes differentially expressed in either of two contrasts (previtellogenic-vitellogenic and vitellogenic-mature); top genes (by mean expression level) in the module; and the most significantly overrepresented Gene Ontology (GO) categories (5% false discovery rate). All genes encoding the pituitary peptide hormones are amongst the highest expressed, and shown in bold. GO overrepresentation was not statistically significant for module 12, which contains few genes (grey text).

Based on the expression of top genes, as well as on unbiased Gene Ontology overrepresentation tests, nearly all of these modules are linked to specific biological processes. In addition, many show a clear association with one or more maturation stages. For example, genes related to cell division are strongly downregulated in adult fish pituitaries (module 19), while protein translation increases in these animals (module 7). The peptide hormone genes showing an increase over maturation are split between two large modules (4 and 6), while those decreasing are grouped in module 18 (figure 3). The consistently highly expressed *prl* is placed in a module (14) which shows no correlation with developmental pseudotime.

### Specific cell types are responsible for distinct developmental expression trends

As it needs to meet changing demands during development, the teleost pituitary gland is known to undergo compositional changes over time^2^. The dynamic landscape of transcriptional trends seen in figures 3 and 4 might therefore reflect changes in cell type composition, in addition to changes in the regulation of gene expression within cells.

We have recently described a single-cell transcriptomics (scRNA-seq) dataset^3^ on the medaka pituitary (figure 5a). These data comprise shallow transcriptomic profiles for 6396 cells (2592 from mature female, 3804 from mature male fish). Clustering of cells by similarity in expression (figure 5a) reveals distinct cell types linked to the production of specific peptide hormones. Several large and small clusters of cells remain functionally uncharacterized (labelled A, B and C in figure 5a).

**Figure 5.**
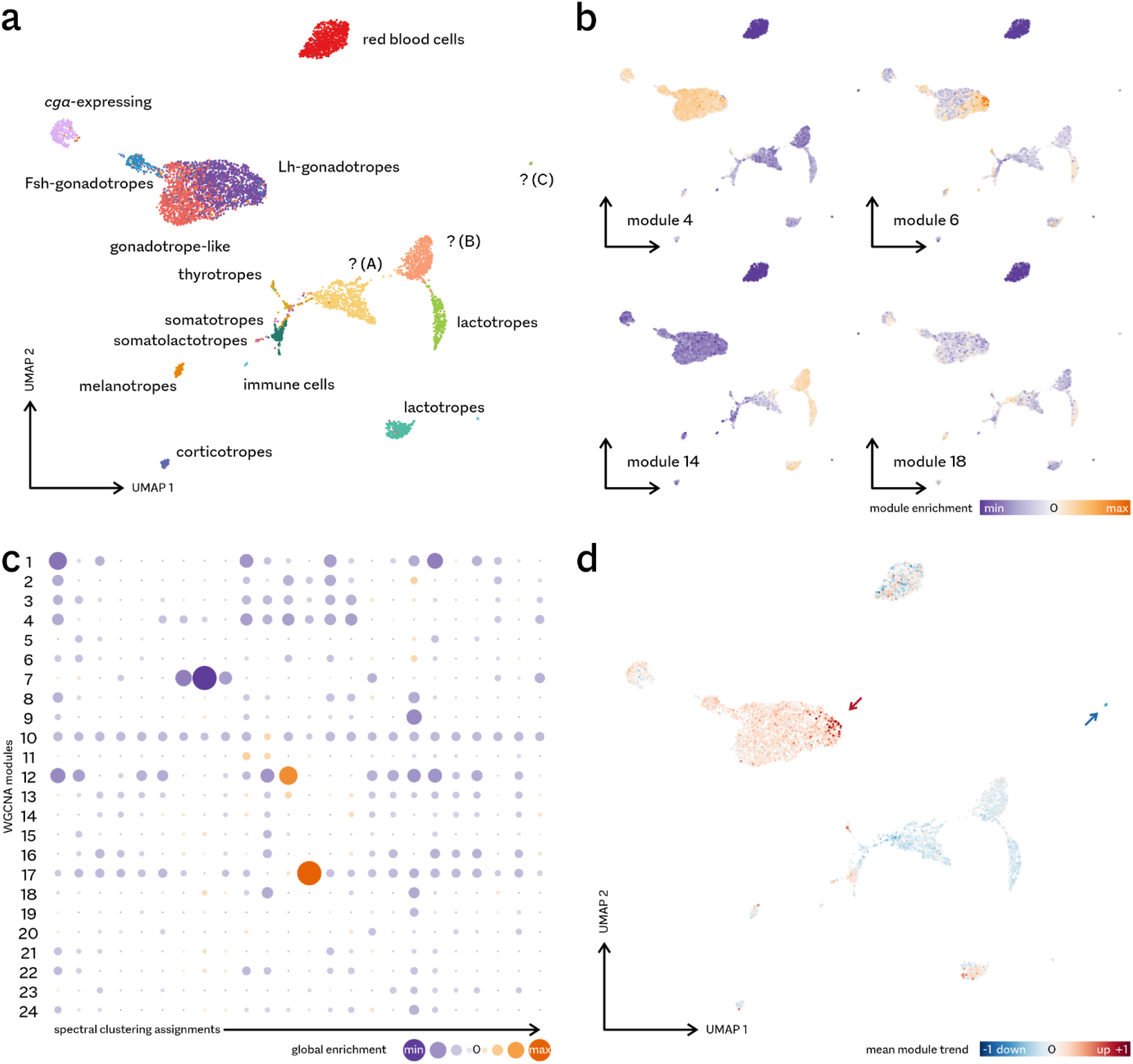
Integration of expression modules with single-cell RNA-seq data. **a** In this UMAP (uniform manifold approximation and projection) visualization of medaka pituitary scRNA-seq data, cells cluster together if their transcriptomic profiles are similar. Based on this, we previously recognized 16 distinct cell types^3^. No function was defined for three of these cell clusters (marked A, B and C). **b** Examples of time series module association with single cells. Genes associated with specific modules (cf. figure 3) exhibit clear concerted under- or overexpression patterns across scRNA-seq clusters. Module 4 and 6 (increasing during development) appear mostly linked to gonadotropes and related cell types; module 14 (stable) is strongly enriched in lactotropes and an uncharacterized cell type, while module 18 (decreasing) shows a relatively broad expression pattern. Dark green (minimal enrichment) typically indicates a module is not expressed at all in a cell. **c** Cell type clustering based on module enrichment scores per cell. Here, a spectral (graph-based) clustering approach was used to assign cells to one of 24 predicted cell types. Resulting cell cluster often show strong enrichment or underrepresentation of specific expression modules. **d** Weighting enrichment scores by PC1 pseudotime (see figure 3) reveals which cells gain or lose prominence over the course of sexual maturation. Arrows indicate cells that are especially strongly positively or negatively correlated with time.

Integrating expression modules (figure 4) with these scRNA-seq data can potentially uncover if certain cells are responsible for specific expression trends. In order to achieve this, we summarized the scRNA-seq expression in 24 enrichment scores per cell, one for each expression module. Briefly, these values are based on the ratio between the observed and expected number of genes per module found expressed in the cell (see Methods for details). As such scores only consider transcript presence or absence, they eliminate the influence the actual expression levels involved.

These enrichment scores expose the investment of a cell in a particular module of genes, relative to the other modules. When visualized per individual cell, scores for a specific module often present a distinct expression pattern over cell types (figure 5b and supplementary figures 4, 5). For example, genes in module 14 are predominantly expressed in lactotropes and one of the uncharacterized cell types, which is consistent with the inclusion of *prl* in this module (see figure 3c). Note that the variation in expression of *prl* itself, the most highly expressed gene in the compound pituitary, contributes only 1.67% to this enrichment, as it is based on a full module comprising 60 genes.

A summary of enrichment scores for all cells of an annotated type often shows clear associations between cell types and expression modules (supplementary figure 6). In fact, the 24 enrichment scores per cell offer sufficient discriminating signal to form the basis for *de novo* cell type clustering (figure 5c and supplementary figure 7; note that this analysis excludes the red blood cells, which have very low transcript counts and as a result often display enrichment artifacts, supplementary figure 8). This module-based cell clustering highlights that modules 12 and 17 are especially strongly associated with specific cell types (figure 5c).

Finally, some cell types show clear enrichment for the modules that change over time, suggesting these cells change either in number or in transcriptional regulation. Figure 5d offers a single, integrated perspective on cellular function in development and sexual maturation. Here, each cell receives a verdict reflecting its relative importance over time (based on weighting the enrichment scores by correlation with the PC1 ordering, see figure 4).

This unbiased data integration reveals that most hormone-producing cell types increase in importance during sexual maturation, with especially the Lh-producing gonadotropes gaining in prominence (red arrow in figure 5d). Intriguingly, the analysis also suggests that all three previously uncharacterized cell types (A–C) lose significance over time, with the strongest decrease associated with one of the cell clusters that is smallest in adult fish (C, blue arrow in figure 5d).

### Rare cell populations ectopically express lipid homeostasis genes

These links between cell types and temporal expression modules could potentially help determine the identity, functions, and developmental dynamics of the three uncharacterized cell types (A, B and C in figure 5a). In particular, small cell cluster C shows the strongest associations in this analysis. It is affiliated with expression modules 12 and 13 (supplementary figures 6 and 9), which show some of the most prominent expression changes over later development (figure 4), with 86% of genes in either cluster differentially expressed between the vitellogenic and fully mature stages.

Uncharacterized cell cluster C comprises only 21 cells, which co-localize in the global UMAP view (figure 5a) with five cells previously classified as belonging to uncharacterized cluster A. By scRNA-seq pseudotime analysis, these 26 cells appear more similar to each other than to the bulk of cluster A (figure 6a). They differ, however, by the expression of modules 12 and 13 (figure 6b), which are mostly specific to the 21 cells (‘group I’ in figure 6). The five cells belonging to cluster A (‘group II’) are instead characterized by enrichment for the expression of module 17 genes (figure 6b).

**Figure 6.**
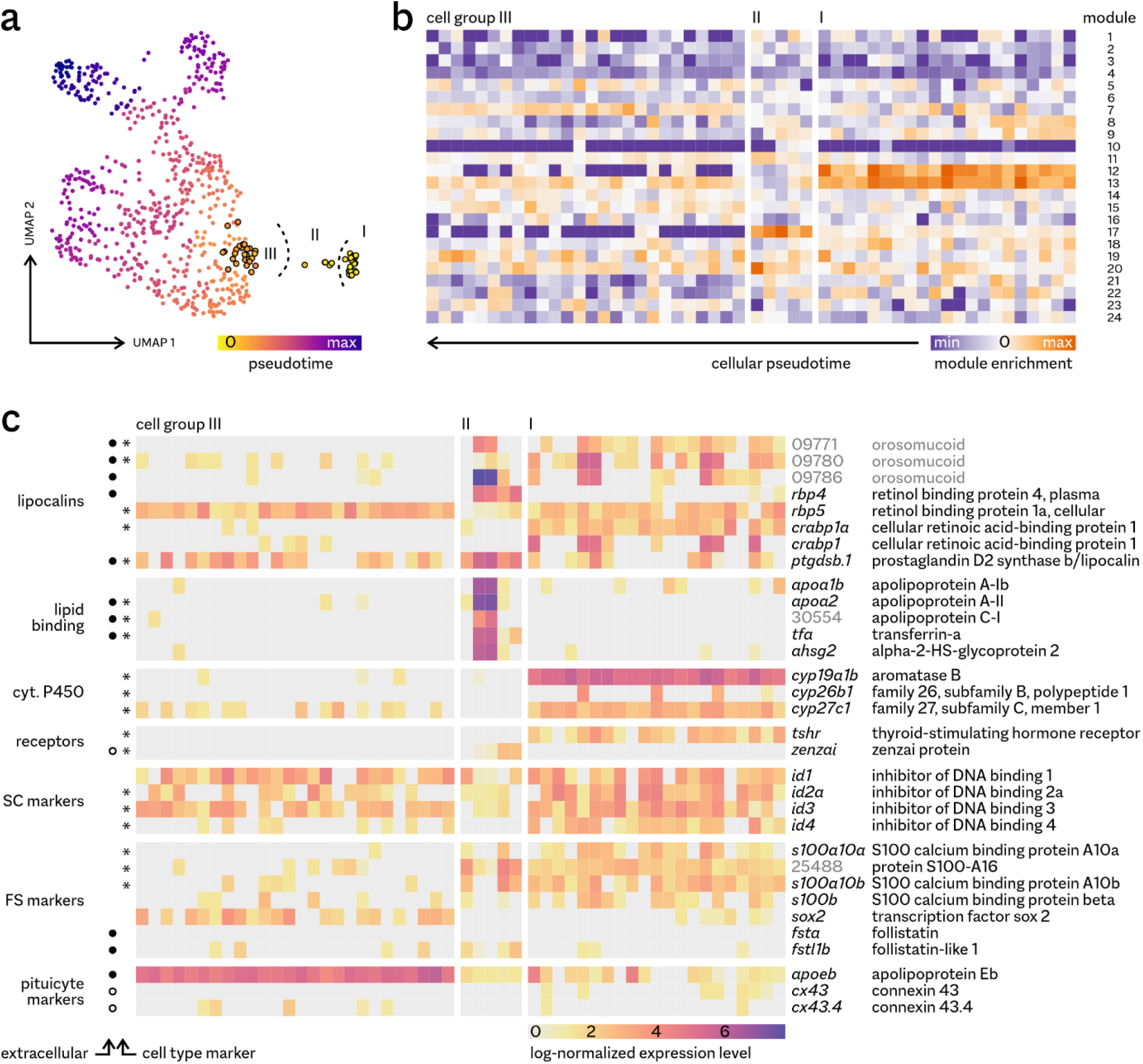
Gene expression profiles of rare cell groups. **a** Definition of cell groups based on cellular pseudotime. In this new UMAP projection focusing on cell clusters A and C only (see figure 5a), cluster C is taken as the root for pseudotime inference (pseudotime 0). Group I corresponds to cluster C; group II to cluster A cells that are close in pseudotime to group I; and group III is a control set shown for contrast, consisting of 26 cells of the main bulk of cluster A closest in pseudotime to the two small cell groups. **b** Expression module enrichment for the 52 cells in these three groups. **c** Selected highly and/or differentially (*) expressed genes suggest functional roles for the rare cell populations. Predicted extracellular protein products are marked by black circles, cell membrane proteins by open circles (supplementary table 3). See supplementary figures 10–15 for expression patterns across all pituitary cell types. SC: stem cells, FS: folliculo-stellate cells.

This module enrichment is also borne out by an analysis of differential expression between rare cell groups I & II and the rest of the pituitary. These cells (together accounting for 0.4% of scRNA-seq expression profiles) are characterized by the differential overexpression of a total of 134 genes. 74 of these are highly expressed in the compound pituitary at some developmental timepoint, and therefore included in the WGCNA analysis. Amongst these 74, genes from the three modules are strongly overrepresented: the set includes 43% of module 12 genes, 26% of module 13, and 45% of module 17.

The marker genes and the genes generally highly expressed in these cells contain clear functional patterns (figure 6c, supplementary figures 10–15). Both groups show high expression of several members of the lipocalin/calycin family: small, usually extracellular proteins with lipophilic molecule binding functions^9,10^. These include retinol/retinoic acid-binding proteins, as well as three unnamed homologues of orosomucoid (alpha-1-acid glycoprotein), which in mammals is a plasma carrier protein of lipophilic compounds, typically expressed in the liver. In addition, the five cells in the smaller group II (less than 0.1% of cells in the adult pituitary) express several other lipid homeostasis and transport genes, including *apoa1b* and *apoa2*, encoding the apolipoproteins A. Two cells show especially high expression levels.

Consistent with these observations, the 21 cells in the larger group I (0.3% of pituitary cells) specifically express cytochrome P450 enzymes involved in the synthesis of lipophilic compounds. *cyp19a1b* encodes aromatase B, the brain enzyme converting androgens to estrogens and involved in steroid feedback along the BPG axis^4^; *cyp26b1* and *cyp27c1* encode enzymes involved in processing retinoids. In addition, both cell groups specifically express putative receptor genes for the lipid transport genes involved (supplementary figure 12). The receptor for thyroid stimulating hormone has been shown to respond directly to human orosomucoid^11^; one of its teleost orthologues^12^, *tshrb*, is expressed exclusively in the 21-cell group I. *zenzai*, specifically expressed in the five cells of group II, is one of five medaka orthologues of scavenging receptor B, which binds apolipoprotein A^13^.

Finally, the differentially expressed genes contain an indication of cell type identity. Several of the retinoid-related genes, as well as aromatase, are expression markers for pituicytes, a population of astroglia-like cells of the pituitary and hypothalamus^14^. In addition, all three uncharacterized clusters express S100 genes (supplementary figure 13), which encode calcium-binding proteins. Four paralogues are specific to the smaller groups (figure 6c). This protein family is the established marker for both pituicytes and pituitary folliculo-stellate cells^15,16^, a heterogeneous population presumed to be involved in paracrine communication^17^ and exhibiting some of the characteristics of stem cells^18–20^, including expression of *sox2*. Interestingly, this possible stem cell-like nature of these cells is supported by the high expression of all four medaka paralogues of the *inhibitor of DNA binding* developmental/stemness transcription factors (supplementary figure 14).

### Folliculo-stellate cell populations decline during maturation

The entire heterogeneous ensemble of folliculo-stellate cells and pituicytes appears to become less prominent in the bulk transcriptome during development (figure 5d). In general, this decrease is relative to other cell types, and may therefore simply be the result of the proliferation or transcriptional activation of hormone-producing cell types. However, the association of small cell groups with strongly declining trends, as seen for the cells expressing lipid homeostasis genes, suggests such cells may actually vanish from the pituitary during sexual maturation.

Modules 12 and 17 are highly specific to these small groups of cells, while module 13 genes are more widely expressed across all three uncharacterized clusters (supplementary figures 4, 5). Especially module 17 genes are exclusively highly expressed in only the five cells of group II (figures 6b, 7a, supplementary figure 11), with the highest expression found in just two cells (figures 6c, 7b). These highly expressed lipid carrier genes (including *apoa1b* and *apoa2*, encoding the apolipoproteins A) are typically specific to the liver, with their protein products secreted into the bloodstream. This homology-based functional annotation is corroborated by computational prediction of the subcellular location for the encoded proteins, which for nearly all of these genes strongly supports a secreted product (figure 6c).

**Figure 7.**
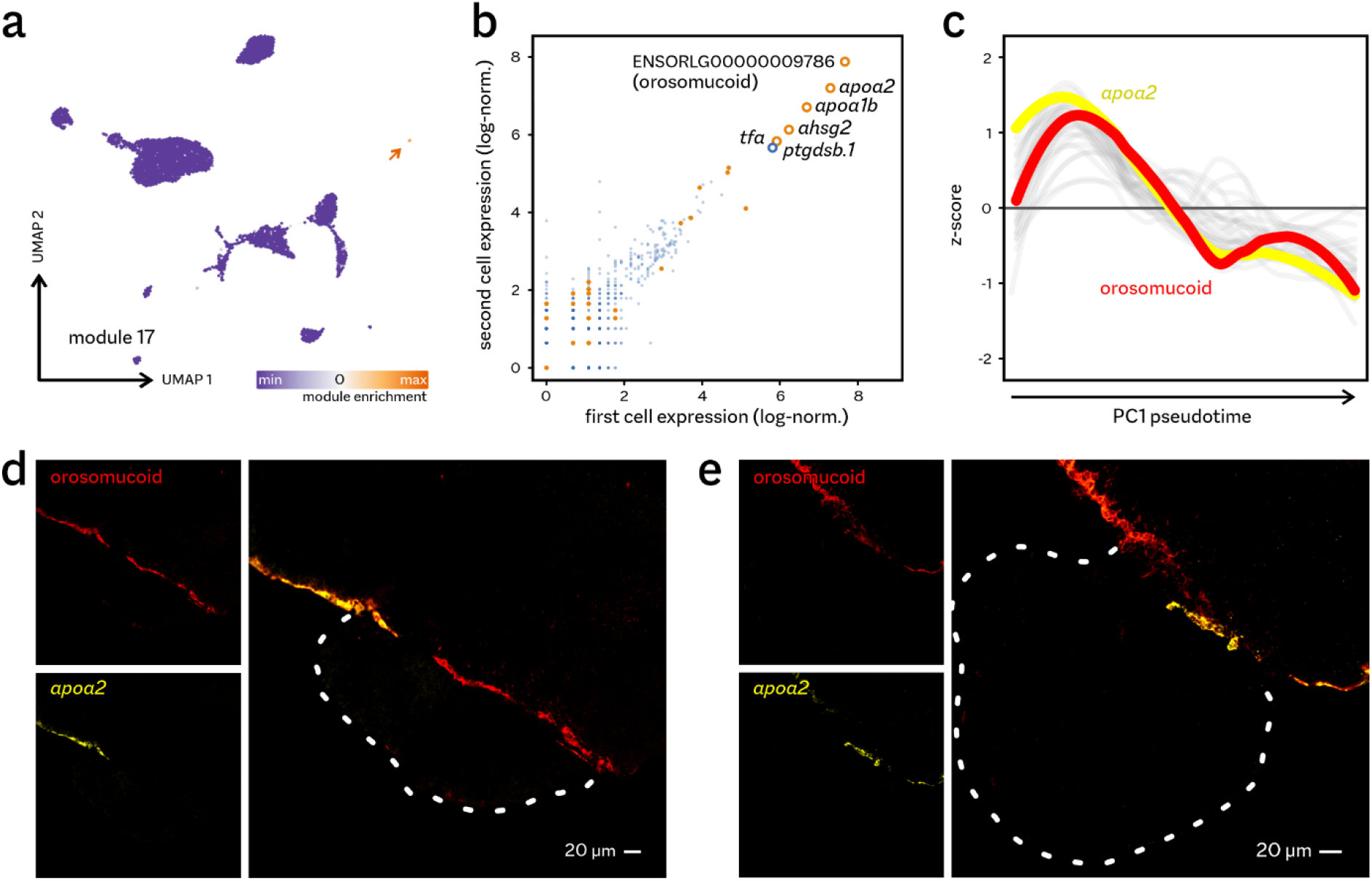
Rare cells display liver-like expression profiles. **a** Expression of module 17 genes is almost entirely limited to the five cells of group II (arrow). **b** Two cells show especially high expression of module 17 genes (orange). **c** Expression of genes in module 17 follows a strong declining trend over development. See supplementary figure 9c for a heatmap visualization for all genes. **d, e** RNAScope *in situ* hybridization on parasagittal sections of the juvenile (two months, **d**) and adult (six months, **e**) female pituitary show expression of module 17 genes along the brain-pituitary interface, with limited expression in cells of the pituitary proper (pituitary gland outlined in white dashed lines).

Expression of these genes has not been detected in the teleost pituitary gland before^21,22^ (supplementary figure 16). In order to exclude the remote possibility of contamination with liver tissue, we compared total gene expression in the adult pituitary to that in liver^23^. Although *apoa1b* and *tfa* are indeed the most abundant transcripts in the medaka liver, overall, the liver and pituitary transcriptomes are incompatible (supplementary figure 17). Highly expressed liver genes are not detectable at substantial levels in the pituitary transcriptomes, while the most highly expressed lipid carrier, orosomucoid ENSORLG00000009786, is barely expressed in the liver at all. Interestingly, this gene is one of three (divergent) local duplicates, of which the other two *are* expressed in both the liver and the pituitary (supplementary figure 17).

The three expression modules associated with the small groups of cells are all characterized by decreased expression in adults (figure 4). In the younger fish of the time series, expression of several module 17 genes is as high as that of the established peptide hormones, declining precipitously in sexually mature animals, with the lowest expression observed in the most mature animals (figure 7c, supplementary figure 2). As rare, specialized cells are responsible for most of the overall expression of module 17 genes (figure 7b), this suggests that this cell type is much more numerous in the juvenile pituitary gland.

We tested this hypothesis using *in situ* hybridization for the two most highly expressed lipid carriers in both juvenile and adult medaka (figure 7d, e). Expression of both *apoa2* and orosomucoid ENSORLG00000009786 coincides in cells located at the boundary of the pituitary and the hypothalamus, with orosomucoid showing wider distributions along the brain’s edge and in the pituitary. In contrast to the RNA-seq data, no major decline in distribution or expression level is apparent. This suggests that the trend seen in the time series may be partially the result of differences in pituitary sampling efficiency for different life stages, although this does not explain the peak in expression after early pre-vitellogenesis (figure 7c).

We subsequently investigated whether the expression pattern of the similar group I of 21 cells fits this explanation (figure 8). Aromatase (*cyp19a1b*) is most strongly expressed in these rare cells, but shows a wider distribution pattern over the pituitary with medium expression in gonadotropes and thyrotropes (figure 8a). The thyroid-stimulating hormone receptor (*tshrb*) is exclusively expressed in group I, with no expression in other pituitary cell types (figure 8b). Orosomucoid ENSORLG00000009786 is expressed in both the rare cell groups, and at low levels in other cell types (figure 8c).

**Figure 8.**
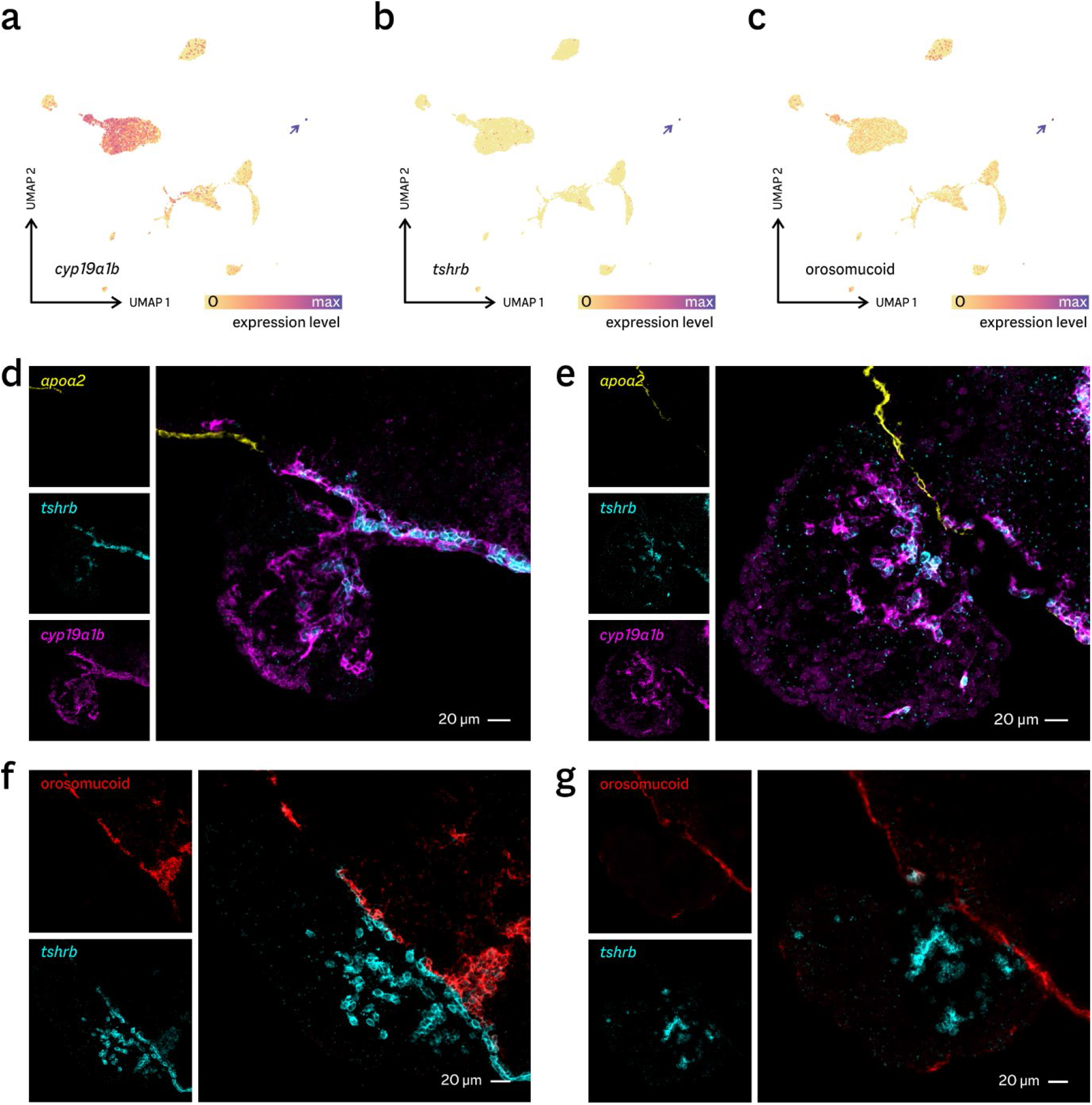
Expression of lipophile-associated genes. **a–c** Expression levels superimposed on the scRNA-seq cells in UMAP view. **a** Aromatase (*cyp19a1b*) is expressed in both gonadotropes and group I cells (arrow), but at much higher levels in the later. **b** The thyroptropin receptor (*tshrb*) is exclusively expressed in group I cells. **c** Orosomucoid (ENSORLG0000009786) is highly expressed in group II cells, with limited expression throughout the pituitary. **d–f** RNAScope *in situ* hybridizations confirm these expression patterns in juvenile (**d, f**) and adult (**e, g**) pituitary glands.

*In situ* hybridization (figure 8d–g) again shows all to be strongly expressed at the brain-pituitary interface, often confirming co-expression as expected (see figure 6c). The interface cell population expressing both *tshrb* and aromatase is distinct from the cells expressing *apoa2* (figure 8d, e), but partially coincides with the expression of orosomucoid (figure 8f, g). In addition, both *tshrb* and aromatase (and, to a lesser degree, orosomucoid) are co-expressed in the centre of the pituitary gland, with *tshrb* expression decreasing in adult animals, consistent with its inclusion in module 12.

These results, therefore, confirm our inference from integrated RNA-seq data on the existence of previously unidentified folliculo-stellate/pituicyte populations and their developmental dynamics. Their strong expression of key interaction partners of several endocrine axes suggests these cells play a major unrecognized role in teleost endocrine physiology.

## Discussion

In this study, we have probed the development of the medaka pituitary gland by combining longitudinal RNA-seq data with in-depth scRNA-seq for a single timepoint (figure 1). The association between many developmental expression trends and cell type clusters (figure 5) allows us to extrapolate the transcriptional activity of cell types over time (figure 5d). This is analogous to deconvolution of each RNA-seq sample to cell types, but without the need for assumptions on the transcriptomic states of the cells in the younger fish. Here, this approach identifies and characterizes novel populations of cells that are transcriptionally prominent in juveniles, yet physically indistinct in adults, without the need to perform scRNA-seq on challenging juvenile samples.

A potential disadvantage of this methodology is that different cell types need to exhibit distinct developmental dynamics and unique expression profiles, and that it requires cell types to be at least detectable at the single scRNA-seq timepoint. In practice, the association between cell types and trends is unlikely to be absolute and exclusive – several large expression modules contain housekeeping genes commonly expressed across many cell types. It does, however, offer an accessible starting point for exploring rich and complex single-cell datasets. The modules derived from time series data can provide a finer-grained classification of cell populations than the clustering based on scRNA-seq data only. For example, the enrichment of modules across the gonadotropes suggests a classification in sub-types that does not entirely conform to the subdivision based on the expression of either *fshb* or *lhb* (supplementary figures 4, 5 and 7).

In particular, this approach will be useful for disentangling the non-endocrine cell populations in the pituitary gland, which show considerable heterogeneity both at the level of scRNA-seq cluster definitions and module enrichment. This diversity is in line with their identification as the transcriptomic counterparts of cytologically defined folliculo-stellate cells^15,16,19^ and pituicytes^14^. The contribution of these cell types to the pituitary transcriptome wanes over the course of sexual maturation (figure 5d). As they both display stem cell-like expression patterns (supplementary figures 14, 15) and are the only cells enriched for cell cycle-related genes (module 19, see supplementary figure 5), this is consistent with a role of folliculo-stellate cells as progenitors for hormone-producing cell types^18–20^

In general, the scRNA-seq profiles agree with previously established transcriptomics markers for zebrafish (*Danio rerio*) pituicytes^14^ and tilapia (*Oreochromis niloticus*) folliculo-stellate cells^24^, including the expression of *sox2* and S100 (supplementary figures 10, 13, 15). However, the expression of markers is not constant over the cell populations (see figure 6c), and no single cell expresses a full set. While this can be partly attributed to species differences, a more reasonable explanation is that previous efforts profiled these heterogeneous populations in bulk (based on the uptake affinity of both pituicytes and folliculo-stellate cells for the fluorescent dipeptide β-Ala-Lys-AMCA). Our integrated dataset therefore forms an important resource for studying the functions and dynamics of distinct cell types within the folliculo-stellate/pituicyte complex.

As an example, we here investigated in detail two small cell populations that appear to be responsible for a major, previously unappreciated contribution to the pituitary transcriptome. These rare cells ectopically express genes not expected in the pituitary, most notably those encoding apolipoprotein A-I and A-II. These proteins are the structural components of high-density lipoprotein (HDL) particles produced and secreted by the liver^25,26^. We confirmed the expression of *apoa2* at the pituitary-brain boundary by *in situ* hybridization (figure 7). In further support of the legitimacy of the phenomenon, we examined recently published zebrafish scRNA-seq data^27^, which confirm the expression of *apoa1, apoa2* and *tfa* in the pituitary gland (supplementary figure 18).

Interestingly, in zebrafish, these genes are expressed in gonadotropes instead of in non-endocrine cells (figure 10). Medaka and zebrafish are only very distantly related, and represent the two largest radiations near the base of the teleost fish clade. The gene regulatory programme(s) responsible are therefore likely common to all teleosts, but they may have been substantially rewired after the divergence of these model species. Together with the expression of different *apoa1* paralogues (*apoa1b* in medaka, *apoa1a* in zebrafish), this opens up the possibility that this programme is an evolutionary novelty ultimately caused by the whole-genome duplication event which shaped teleost evolution and development^28^.

Further evidence on the expression patterns of *apoa1*/*2* in teleosts is inconclusive. In adult carp (*Cyprinus carpio*, a close relative of zebrafish), both genes show a tissue distribution that closely resembles the established mammalian pattern^22^, with no expression at all in the pituitary gland. In several teleosts, however, considerable expression is sometimes detected in the skin, brain, and testis, amongst other tissues^29,30^. In these cases, apolipoprotein A is often assumed to have a local anti-microbial function, a role that is difficult to reconcile with very high expression in and near the pituitary.

The established physiological role of apolipoprotein A is reverse cholesterol transport^26^. HDL particles sequester cholesterol from the bloodstream and deliver it back to the liver. In addition, the gonads and adrenal glands/interrenal tissue interact with HDL to obtain cholesterol, which they need as the raw material for the synthesis of steroid hormones. It is unlikely the pituitary-derived apolipoproteins primarily engage in this scavenging function, as even high juvenile production will be fully eclipsed by the much larger liver. However, many lipoproteins are thought to be involved in delivering messenger molecules in the brain^31^. This suggests the existence of a novel endocrine communication channel along the BPG and BPI axes, in which pituitary-derived HDL-like particles could deliver steroids or other lipophilic compounds to peripheral organs expressing the HDL receptor.

Curiously, one medaka homologue of the HDL receptor (*zenzai*) is co-expressed specifically in the apolipoprotein A-producing cells (figure 6c). In medaka, *zenzai* has been shown to be essential for germ cell survival and gonad development^32^. A similar situation – co-expression of a lipid transport gene and its putative receptor – occurs in the other novel cell type. Both small cell groups express several orosomucoid genes^33^, which in mammals affect thyroid hormone homeostasis through a direct interaction with the thyroid stimulating hormone (Tsh) receptor^11^. In our data, Tsh itself is produced by pituitary thyrotropes (see figure 5a), and the Tsh receptor is exclusively expressed in the rare cells (figure 8). In the human pituitary, the Tsh receptor is expressed in a subset of folliculo-stellate cells^34^.

These observations suggest both rare cell types engage in paracrine (or autocrine) signalling: communication to local cells (or feedback to the producing cell itself) mediated by secreted proteins^17^, a role that is consistent with their identification as folliculo-stellate cell subpopulations^15^. The very high expression levels in the juvenile pituitary, however, do not support an exclusive intra-pituitary paracrine role. *apoa1b, apoa2, tfa*, and orosomucoid ENSORLG00000009786 are amongst the most highly expressed in immature fish, attaining the same levels as the genes encoding the established peptide hormones (supplementary figure 2). Our combined evidence therefore implies teleosts co-opted these secreted liver proteins as endocrine messengers.

## Methods

### Fish husbandry and growth experiment

We studied the pituitary glands and gonads of Japanese medaka (*Oryzias latipes*) of the d-rR genetic background. In addition, these fish express green fluorescent protein under the control of the *lhb* promotor^35^, facilitating pituitary dissection even in small animals. We sampled eggs from a tank containing three females and three males. After hatching in petri dishes, fish from a single egg collection date were transferred to 1-liter tanks in a re-circulation system. After one month, fish were transferred to 3-liter tanks containing exactly 10 fish of the same age. Fish were kept at 28 °C on a 14/10h light/dark cycle, and were fed three times a day with a mix of dry food and *Artemia*. The animal experiments performed for this study followed EU/Norwegian regulations for animal welfare and for the use of animals in experiments.

### Fish sampling

We sampled 118 female fish from our time series experiment (aged 31–178 days after egg collection), as well as the three female parents (375 days) and three mature females of intermediate age (249– 277 days), housed in the same facility. The fish were sacrificed between 09.00–11.00h in the morning by hypothermic shock (immersion in ice water) to minimize distress^36^, followed by severing the spinal cord, and immediate dissection of the pituitary. Pituitaries were carefully dissected using fine forceps and extra care was taken to prevent cross-contamination between samples by carefully cleaning all equipment with ethanol after dissecting each fish, and also before sampling the pituitary. Each pituitary gland was put in an 0.5 ml Eppendorf tube containing 10 μl ice-cold medaka-adjusted^37^ PBS (pH 7.75 and mOsm 290 mOsm) and 5 ceramic beads of 1.4 mm (MP Biomedials), directly followed by the addition of 300 μl Trizol Reagent (Invitrogen) to each tube. The sample was vortexed at high speed for 1 minute and snap frozen in liquid nitrogen. Samples were stored at -80 °C until RNA isolation.

Subsequently, we collected ovaries, transferred them to small glass bottles containing 4% glutaraldehyde (Merck Millipore) 0.1 M phosphate-buffered solution (pH 7.2), and left them to incubate overnight at 4 °C. The samples were then stored in 70% EtOH at 4 °C until histological examination. Whenever possible, ovaries were dissected out and examined macroscopically to obtain a preliminary maturity estimate. Fin clips were collected for sex genotyping.

### Sex genotyping

Genomic DNA was isolated from fin clips by incubation in 25 μl alkaline lysis buffer (25 mM NaOH, 0.2 mM EDTA) for 5 minutes at 95 °C and subsequent neutralization using 25 μl 40 mM Tris-HCl (pH 8.0). Genotypic sex was determined by genomic PCR for the presence or absence of the *dmy* gene (primers 5ʹ-CCGGGTGCCCAAGTGCTCCCGCTG and 5ʹ-GATCGTCCCTCCACAGAGAAGAGA) and analysis by agarose gel electrophoresis^38,39^.

### Transcriptome sequencing

RNA was isolated using a standard Trizol protocol, with minor modifications. Samples were thawed and vortexed for 10 seconds with 120 μl chloroform, incubated at room temperature for 3 minutes, and centrifuged at 14000g for 15 minutes at 4 °C. Glycoblue (Invitrogen) (17.5 μg/ml) was added to each tube to enhance precipitation and visualize the pellet. The aqueous phase (ca. 160 μl) was transferred to a new tube and RNA was precipitated in 160 μl 100% isopropanol by incubation at -20 °C for 1 hour, followed by centrifugation at 14000 g for 15 minutes at 4 °C. The pellet was then washed in 300 μl 75 % ice-cold ethanol and the sample centrifuged at 10000g for 10 minutes at 4 °C. The supernatant was removed and the RNA pellet briefly dried (approximately 5 minutes) at room temperature. The pellet was dissolved in 10 μl of RNase-free water. To avoid unnecessary freeze-thawing of the samples, aliquots of 2 μl of each sample were made for Bioanalyzer analysis. Samples were stored at -80 °C until use.

We investigated RNA concentration and integrity of all samples using the Bioanalyzer Eucaryote total RNA pico kit. All samples included in the downstream RNA-seq analysis had RIN values between 7–10 (mean 9.3, median 9.6).

We selected 68 samples for sequencing, of which 54 are included in the present analysis (13 additional sequenced pituitaries were from fish selected as growth curve outliers during the experiment; one sample was an outlier after sequencing). RNA was shipped on dry ice for library preparation and sequencing at Future Genomics Technologies (Leiden, The Netherlands). cDNA was generated and amplified using the Smart-SEQ HT Kit for low RNA input according to the manufacturer’s recommendations (Takara Bio), which was used for generating sequencing libraries with the Nextera XT kit. Libraries were sequenced at 151 nt paired-end on an Illumina NovaSeq 6000 system.

### Histological analysis

The fixated tissue was dehydrated in a series of increasing concentrations of EtOH (70–100%), each step lasting at least >30 minutes. The last step (100%) was repeated three times and then replaced with approximately 5 ml of preparation solution (100 ml Technovit 7100 with 1 g of Hardener I, Heraeus Kulzer) and kept at slow shaking at room temperature overnight. After infiltration, tissue samples were embedded in cold Histoform S (Heraeus Kulzer) with approximately 1 ml preparation solution and 50 μl Hardener II (Heraeus Kulzer) and incubated at 37 °C. Cured samples were mounted on Histoblocs using Technovit 3040 (both from Heraeus Kulzer). The gonads were sectioned using a Leica Biosystems RM2245 microtome. Sagital sections (3 μm) were made from the periphery until the middle of the gonad and collected on microscope slides every 30–90 μm, depending on the gross maturity of the gonad. Dried sections were stained with Toluidine Blue O (Sigma-Aldrich) and mounted with Coverquick 4000 (VWR International) prior to microscopy analysis. The developmental stage of the tissue was determined by the most advanced ovarian follicle present (see table 1)^7^.

### In situ *hybridization*

We used multiplexed RNAscope *in situ* hybridization to visualize tissue localization of mRNA expression for selected genes: *apoa2* (ENSORLG00000024993), orosomucoid (ENSORLG00000009786), *cyp19a1b* (ENSORLG05548) and *tshrb* (ENSORLG00000014222). Juvenile (two months old) and adult (six monhts old) female medaka were euthanized with an overdose of anesthetic (MS222, Sigma). Adult fish were cardiacally perfused with a solution of 4% paraformaldehyde (PFA) diluted in phosphate buffered saline (PBS) solution (Electron Microscopy Sciences) to remove autofluorescence from blood cells^40^. Brain and pituitary from both adult and juvenile fish were collected and fixed overnight in a fresh solution of 4% PFA at 4°C. Tissues were then rinsed in PBS with 0.01% Tween (PBST, Sigma) before being dehydrated with a series of ethanol solutions of increasing concentration (25, 50, 75 and 96%). Tissues were then stored in methanol for 1 to 6 weeks. Following rehydratation and rinsing in PBS, tissues were incubated in 25% sucrose solution (diluted in PBS) overnight at 4°C, embedded in Tissue-Tek OCT-Compound (Sakura Fintek), quickly frozen over dry ice, and stored at -80 °C. 10 μm parasagittal section were then made with a cryostat (Leica) on frosted microscope glass slides. RNAscope *in situ* hybridization^41^, multiplex version 2, was performed as instructed by Advanced Cell Diagnostics using opal 520, 570 and 690 labels. Confocal plan images were then taken with a Zeiss confocal (LSM 810) with a 25× objective and channels acquired sequentially to prevent cross-signalling of fluorophores. Images were then processed in Fiji^42^ and compiled using Adobe Indesign.

### Additional data

Liver RNA-seq data for two female adult medaka were downloaded from the NCBI Sequence Read Archive (BioProject PRJEB37848, accessions ERS4513972 and ERS4513973). These fish (of the d-rR genetic background) were grown in the same facility as the fish of our main experiment^23^. The data are based on RNA-seq libraries prepared using the Illumina TruSeq Stranded mRNA HT Sample Prep Kit and sequenced at 125 nt paired-end on an Illumina HiSeq 2500 system.

Single-cell RNA-seq expression matrices were downloaded from NCBI Gene Expression Omnibus for medaka (GSE162787) and zebrafish (GSE148591). In the case of medaka, these fish are of the same transgenic line as those used in our main experiment.

### Sequencing data analysis

All FASTQ data were aligned against the medaka HdrR reference genome (Ensembl version 94) using STAR^43^ (version 2.7.6a). Alignments were inspected or processed using Samtools^44^ (version 1.10) and quantified using htseq-count^45^ (version 0.11.2) using the *intersection-nonempty* setting. The resulting data were analyzed in R (version 4.0.3) using the edgeR^46^ (version 3.32.0) and cqn^47^ (version 1.36.0) packages, using the Corrplot (version 0.84) package for visualization. As our data are both amplified for low input material and derive from developmentally heterogeneous samples, they do not fit the assumptions for normalization by scaling commonly applied to RNA-seq (e.g. minimal changes in gene expression between samples). We therefore normalized counts for 23635 protein-encoding genes between samples using quantile normalization as previously described^48^.

Differential expression between maturity cohorts, and along age, length or principal components of the dataset were calculated using edgeR using the GLM functionality. We identified expression trend modules using weighted gene correlation network analysis^8^ (WGCNA package, version 1.69) using default settings (signed network, threshold power 10, and cluster merging at correlation >0.9). Several independent runs with small adjustments in the settings yielded very similar results. Gene Ontology (GO) overrepresentation was analyzed using GOseq^49^ (version 1.42.0) using GO annotations obtained from Ensembl BioMart. We only tested overrepresentation for categories containing more than one gene and with at least one gene in the test group. Initial *p*-values for both differential expression and GO overrepresentation were corrected using the Benjamini-Hochberg procedure to control the false discovery rate at 5%.

Single-cell sequencing data were analyzed using Seurat^50^ (version 3.2.3) for dataset integration and dimensionality reduction. Differential expression between groups of cells was determined using the Seurat *FindMarkers* function (using the default Wilcoxon rank-sum test), using an adjusted *p*-value threshold of 0.05 (Bonferroni correction). The reported differential expression results are the combination of four contrasts: overexpression of group I, II, or both, compared against all pituitary cell excluding the uncharacterized clusters A and C; and differential expression between groups I and II Only genes expressed in at least two cells in the rare groups were considered. Monocle3 (version 0.2.3) was used to infer cellular pseudotime^51^.

Subcellular protein sorting was predicted using SignalP^52^ (version 5.0) and DeepLoc^53^ (version 1.0), available at *www.cbs.dtu.dk/services*.

### Data integration strategy

We used R to combine the time series RNA-seq with the single-cell data by considering over- or underrepresentation of detected module genes within each cell, yielding 24 enrichment scores per cell. As a threshold for detection of expression, to reduce noise, we took at least two sequencing reads. For each module, we calculated the ratio between observed and expected numbers of genes detected:

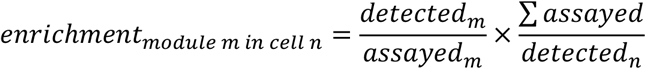

where ‘assayed’ means the numbers of genes involved in, and derived from, the original WGCNA analysis. Finally, to make enrichment scores comparable across different modules, for each module, we centred values by subtracting the module mean and subsequent division by the module standard deviation.

In order to classify cell types based on enrichment scores, we used spectral clustering^54^ implemented in the R package Spectrum (version 1.1). A single weighted integration score per cell was calculated by multiplying each enrichment score by the Spearman rank correlation with PC1 pseudotime (figure 4) for that module, and taking the mean over the 24 numbers.

## Data availability

Raw sequencing files as well as read counts for the time series experiment are available from the NCBI Gene Expression Omnibus (*https://www.ncbi.nlm.nih.gov/geo/*), accession number GSE179598. scRNA-seq data (both reads and Seurat-compatible matrix files) are available through GEO accession number GSE162787.

## Acknowledgements

We would like to thank Lourdes Carreon G. Tan for fish facility maintenance, Simen Sandve for information on the liver data, and Ron Dirks, Kjetil Hodne, and Erik Burgerhout for discussion. This work was supported by the Norwegian University of Life Sciences and the Norwegian Research Council grants no 251307, 255601, and 248828.

## Author contributions

GM, EA-W, F-AW and CH conceived and designed the study. GM and EA-W performed the experiments. KvK analyzed gonad samples. RN-L performed genotyping. RN-L and RF performed *in situ* hybridization. KS contributed data analysis. CH analyzed the data and wrote the paper, with contributions from the other authors.

