## Supplementary figures 1-18 for "Integrative transcriptomics reveals ectopic lipid homeostasis mechanisms in non-endocrine cells of the teleost pituitary"

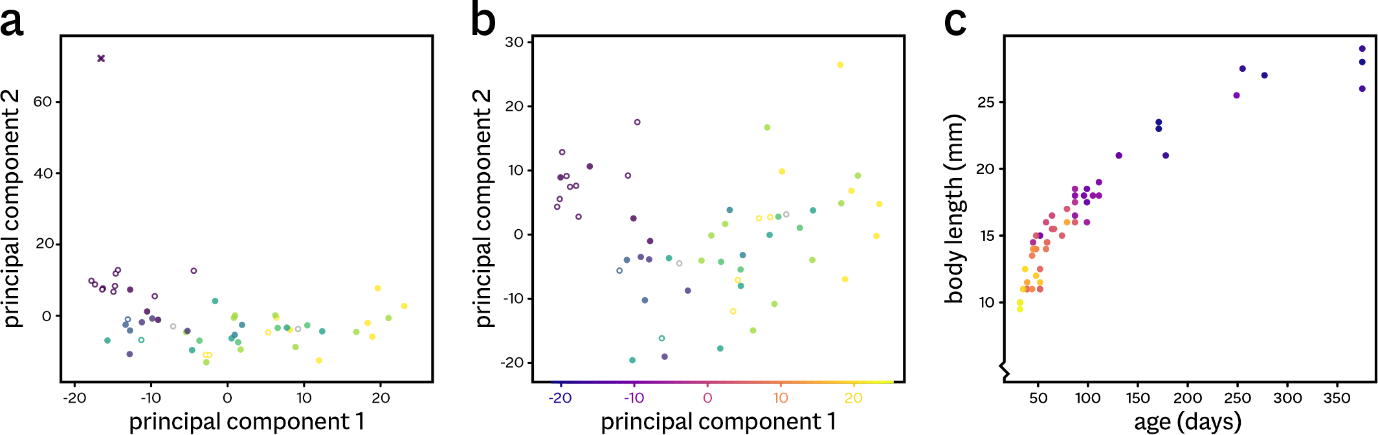

**Supplementary figure 1. Selection and ordering of pituitary samples. a** In an initial principal component analysis (PCA) based on 55 sequenced pituitary glands, one (adult) sample is a clear outlier (cross). We excluded this sample from further analyses. **b** A PCA using 54 samples shows a clear correlation between gonad maturity (sample colour code: see main figure 2) and principal component 1. **c** Sample length and age colour-coded by principal component 1 (see panel b).

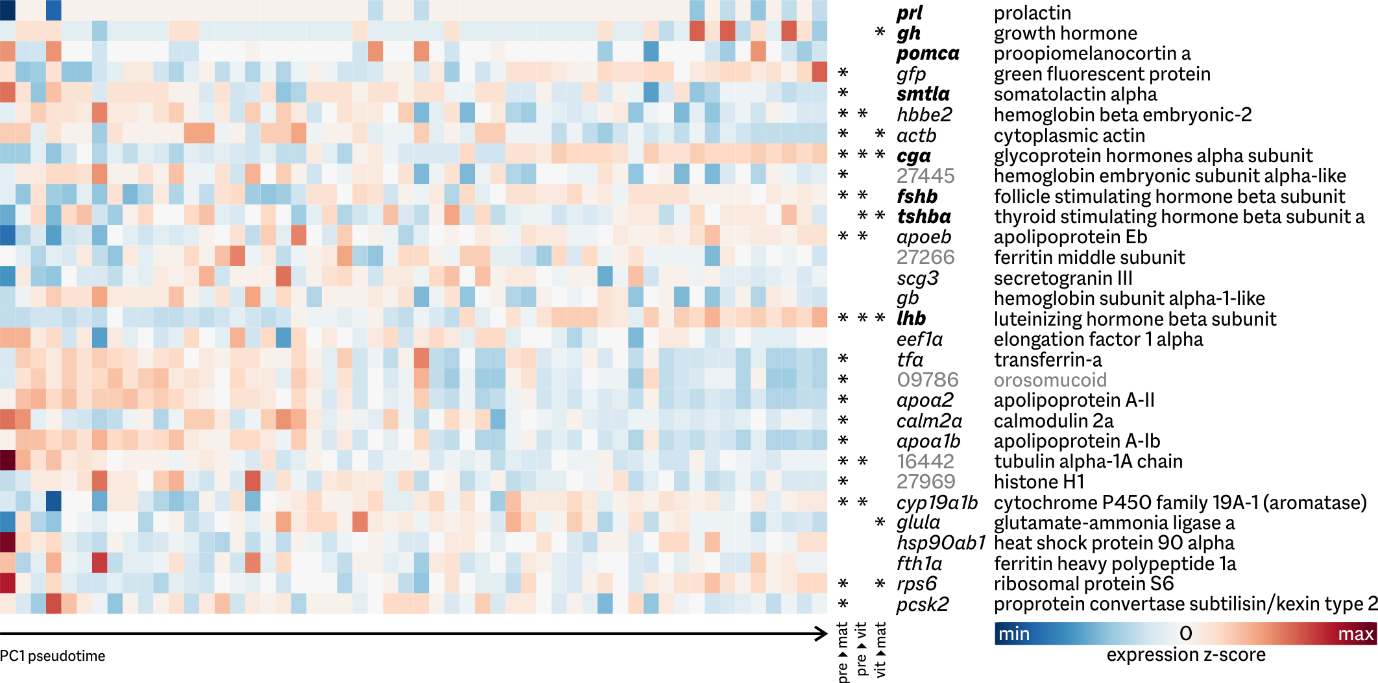

**Supplementary figure 2. Gene expression patterns over development.** Heatmap of expression patterns of highly expressed genes, defined as all genes that are in the top 10 by expression level in at least one sample. Expression values displayed as z-scores (standard deviations around the mean) per gene, scaled to the overall maximum and minimum values, and ordered by pseudotime (see supplementary figure 1). Asterisks indicate significant differential expression (5% false discovery rate) for three contrasts between groups: previtellogenic-mature, previtellogenic-vitellogenic, and vitellogenic-mature (see table 1 for groups).

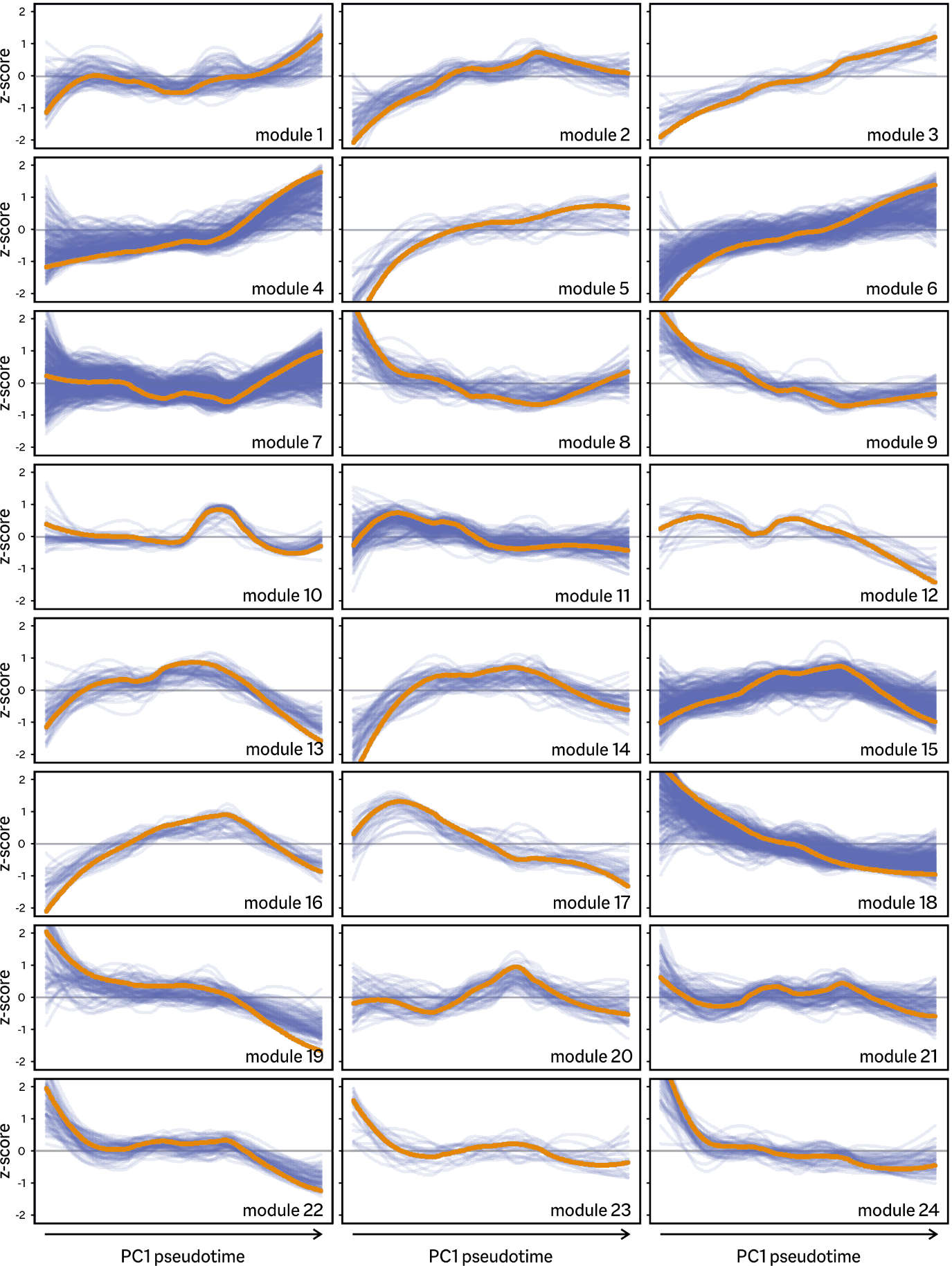

**Supplementary figure 3. Expression trends per WGCNA module.** Gene expression is shown in z-scores (standard deviations around the mean) over development. Blue lines are loess local regression curves per gene, orange curves represent module eigengenes.

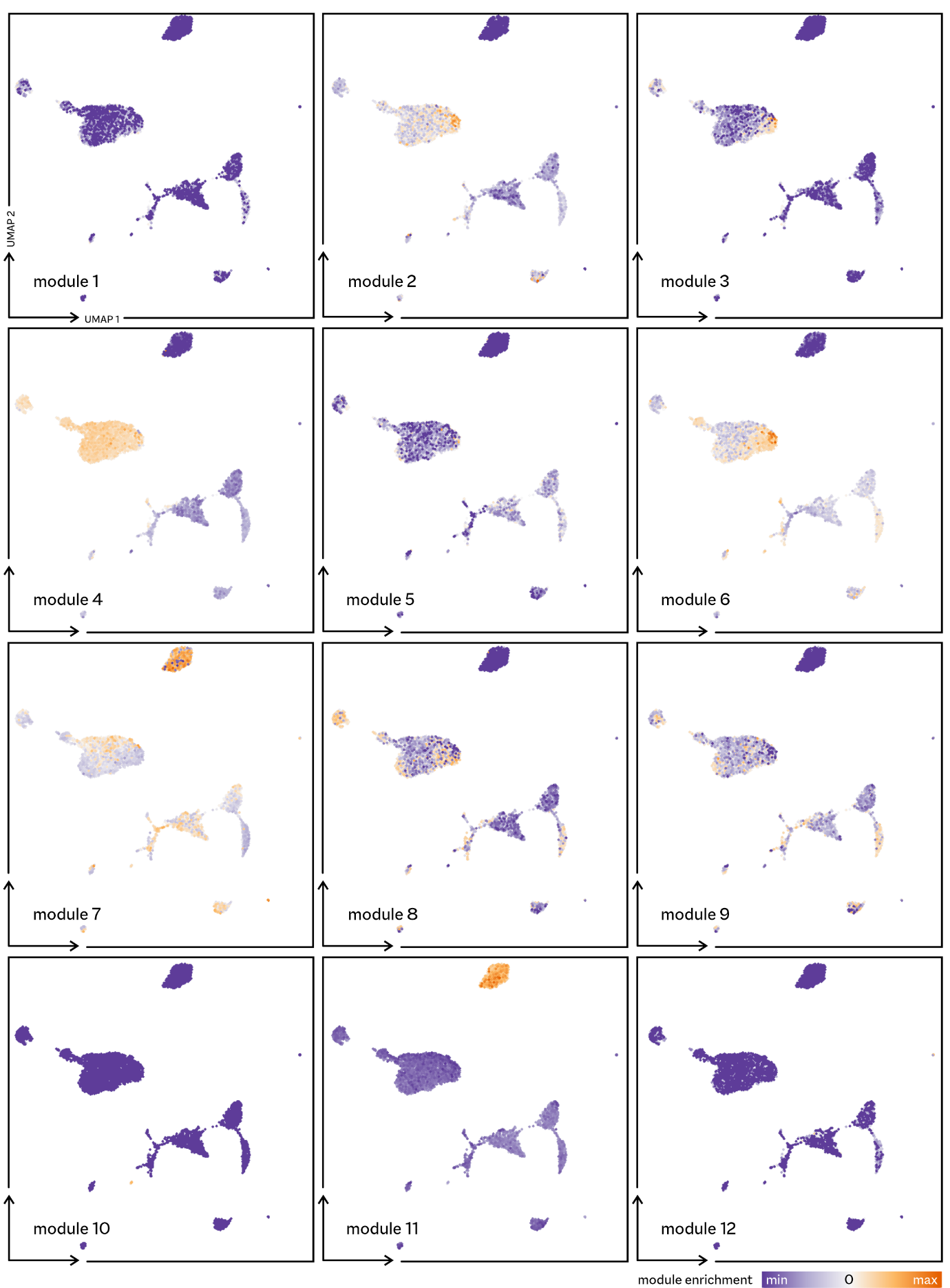

**Supplementary figure 4. WGCNA module enrichment often shows a distinct distribution pattern over cell types**. For cell type annotation in this UMAP projection, see main figure 5.

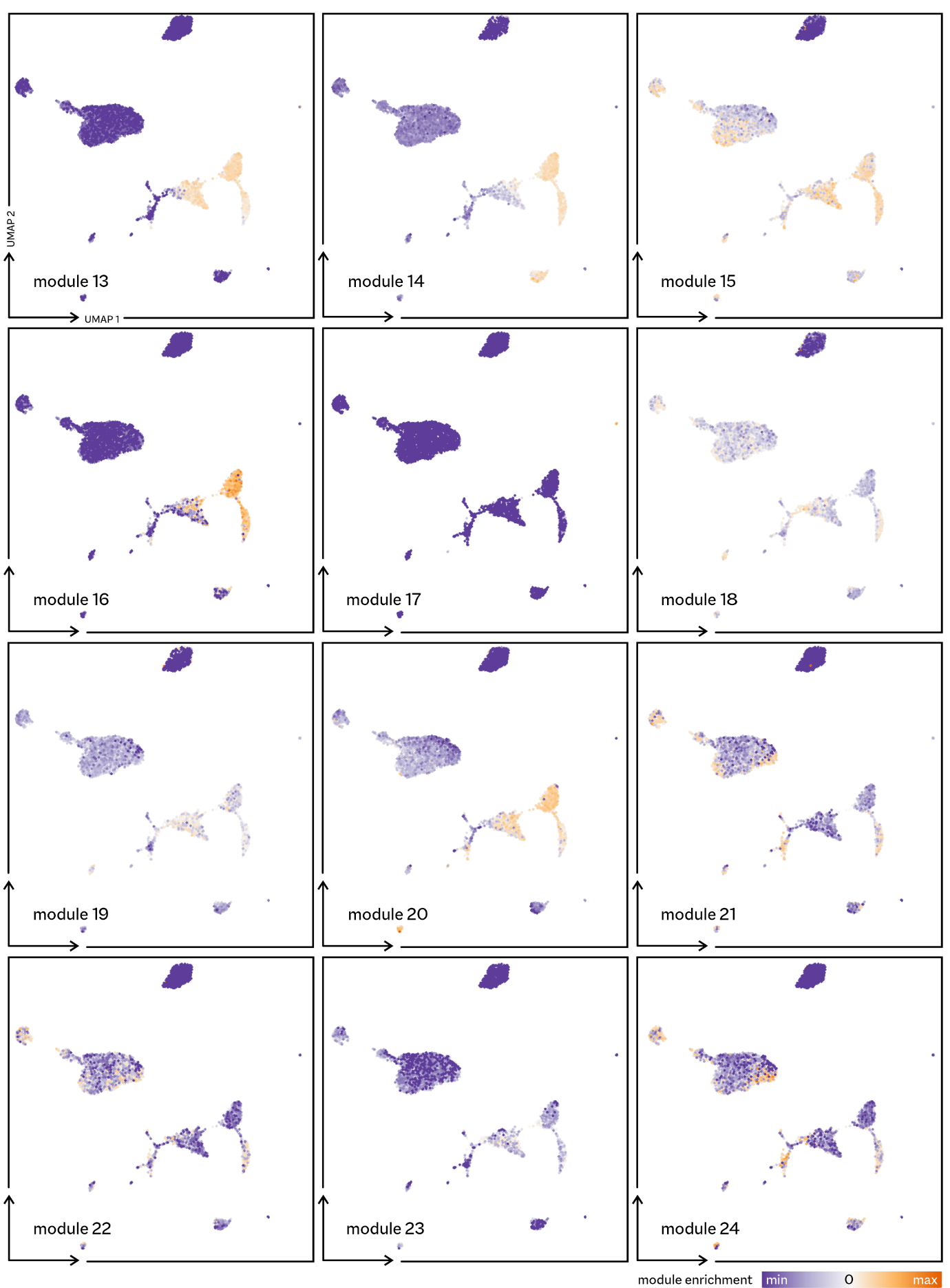

**Supplementary figure 5. WGCNA module enrichment often shows a distinct distribution pattern over cell types**. For cell type annotation in this UMAP projection, see main figure 5.

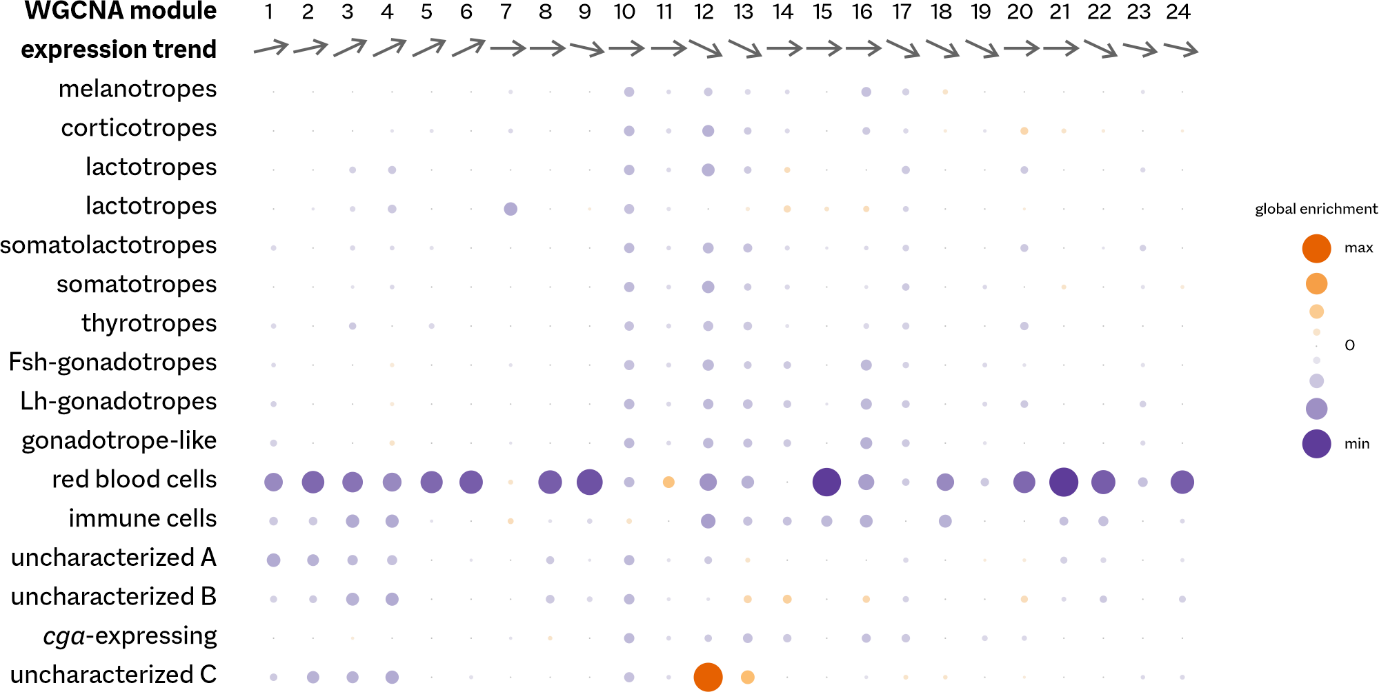

**Supplementary figure 6. WGCNA module enrichment summary per cell type**. Enrichment values per module are scaled globally, showing the strongest associations between modules and cell types over the entire dataset. These enrichment values are affected by annotated cell cluster size, leading for example to strong enrichment of module 12 genes in the small and homogeneous cluster ‘C’ of uncharacterized cells. Red blood cells show strong underrepresentation of nearly all modules, with the exception of module 11 containing hemoglobins. Expression trends are derived from main figure 4.

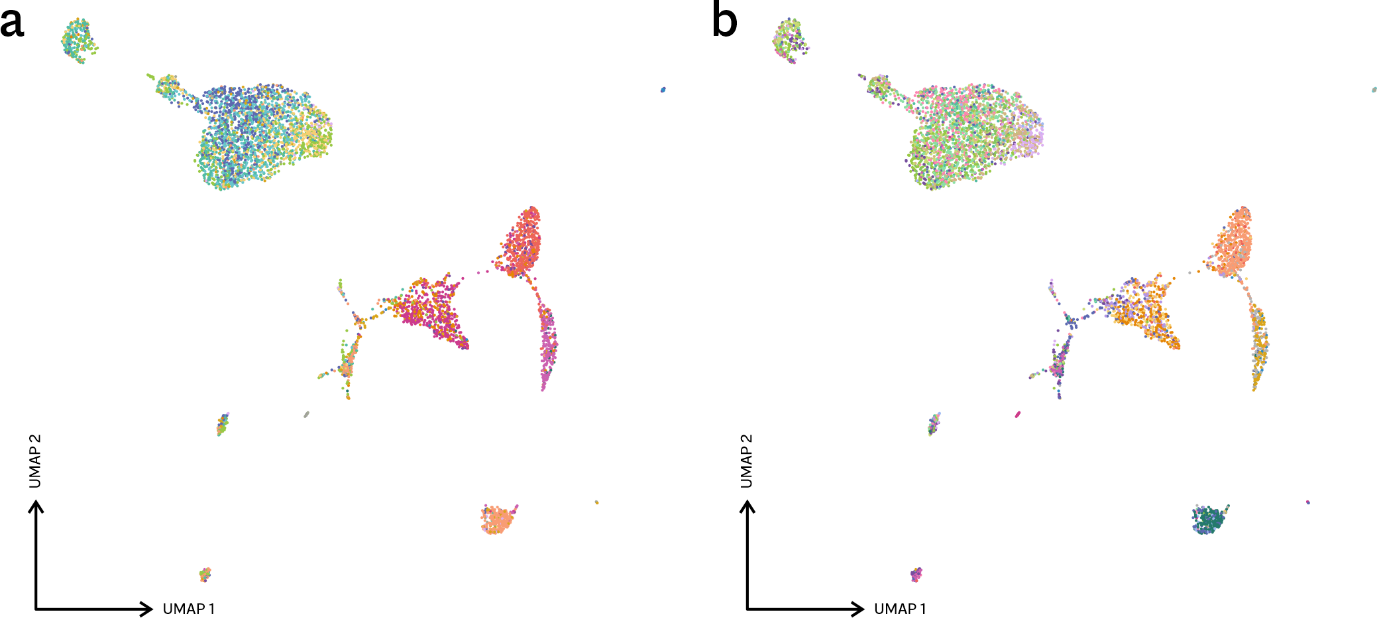

**Supplementary figure 7. Cell type classification by spectral clustering on module enrichment scores.** Colour coding indicates cell type assignments based on either 16 (**a**) or 24 (**b**) spectral clusters. Cells previously identified as red blood cells are not included in this analysis.

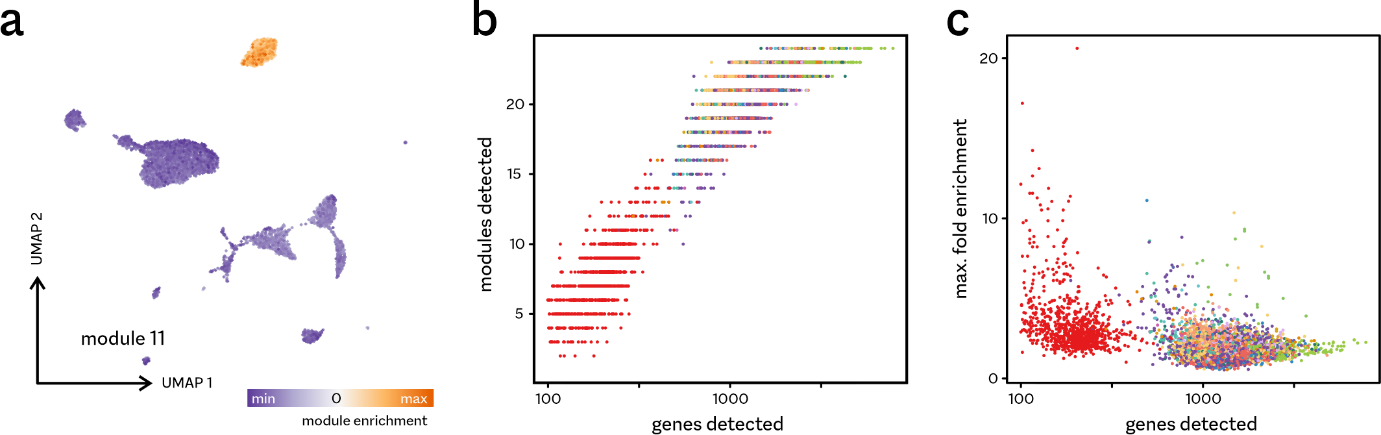

**Supplementary figure 8. Noisy module enrichment in red blood cells**. **a** Module 11, containing predominantly hemoglobin genes, is strongly enriched in red blood cells only. **b** In the scRNA-seq data, these cells are represented by fewer reads, and therefore by fewer expressed genes and detected expression modules. Colours correspond to annotated cell types (main figure 5a), with red dots representing the red blood cells. **c** Lower numbers of detected genes/modules result in artifactually strong enrichment values, represented here by the maximum enrichment per cell over all 24 modules).

**Supplementary figure 9 (*next page*). Expression patterns over development in selected modules.** Heatmaps of expression of all genes in modules 12 (**a**), 13 (**b**) and 17 (**c**). Expression values displayed as z-scores (standard deviations around the mean) per gene, scaled to the maximum and minimum values per module, and ordered by pseudotime (see supplementary figure 1). Asterisks indicate significant differential expression (5% false discovery rate) for three contrasts between groups: previtellogenic-mature, previtellogenic-vitellogenic, and vitellogenic-mature (see table 1 for groups).

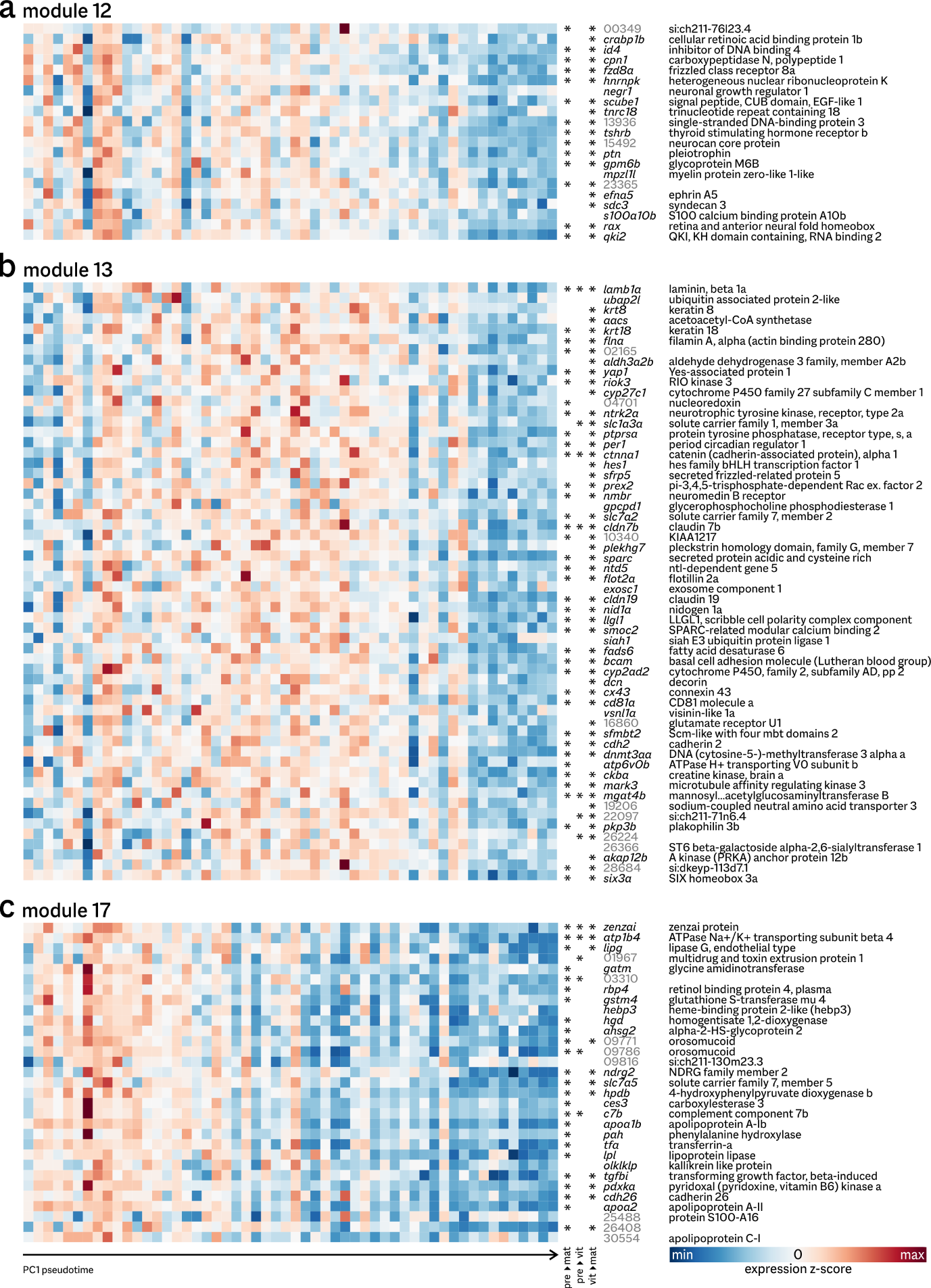

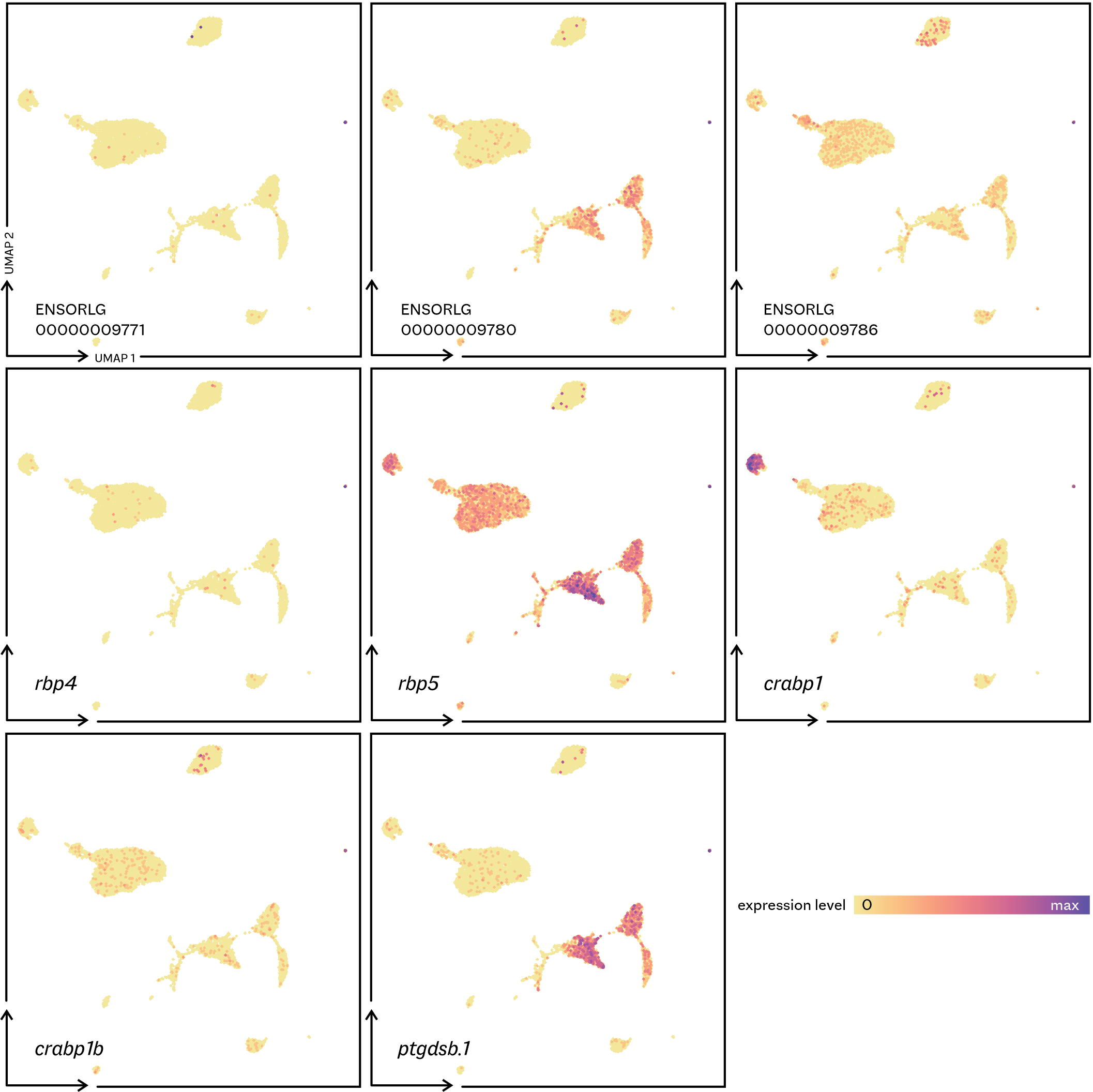

**Supplementary figure 10. Expression of selected lipocalin genes in the pituitary gland.** All are highly expressed in the rare cell clusters (extreme right); some show additional high expression in other putative folliculo-stellate cells and gonadotropes. Occasional strong expression in red blood cells (top cluster) is based on very few reads; as these cells are transcriptionally less active, data normalization scales single reads (presumably derived from ambient RNA) to relatively high expression values.

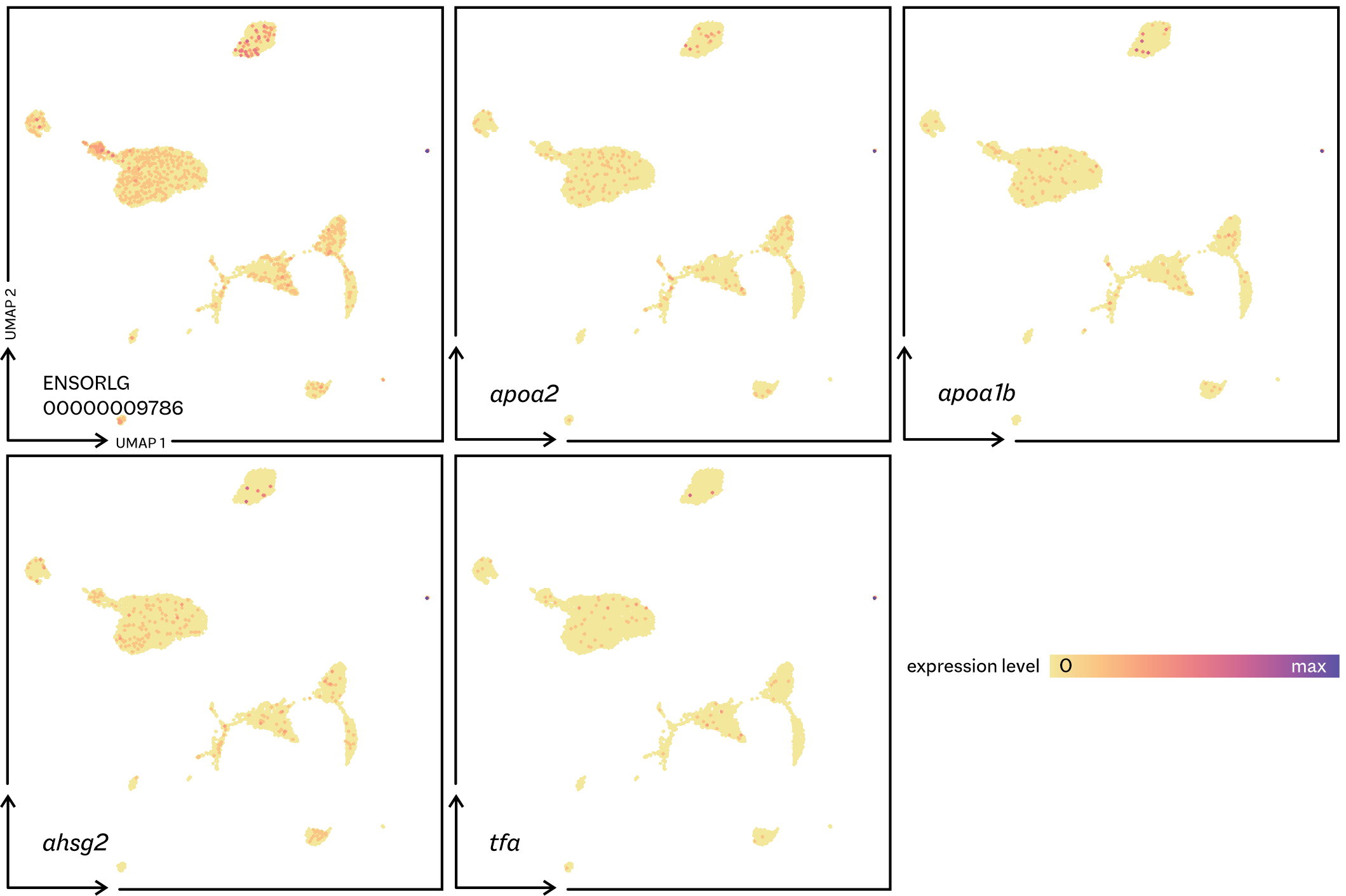

**Supplementary figure 11. Localization of highly expressed module 17 genes to rare cells.** Most of these genes are only expressed in a few cells (small isolated cluster to the extreme right). Only ENSORLG00000009786 (encoding orosomucoid) shows very modest expression in other pituitary cell types.

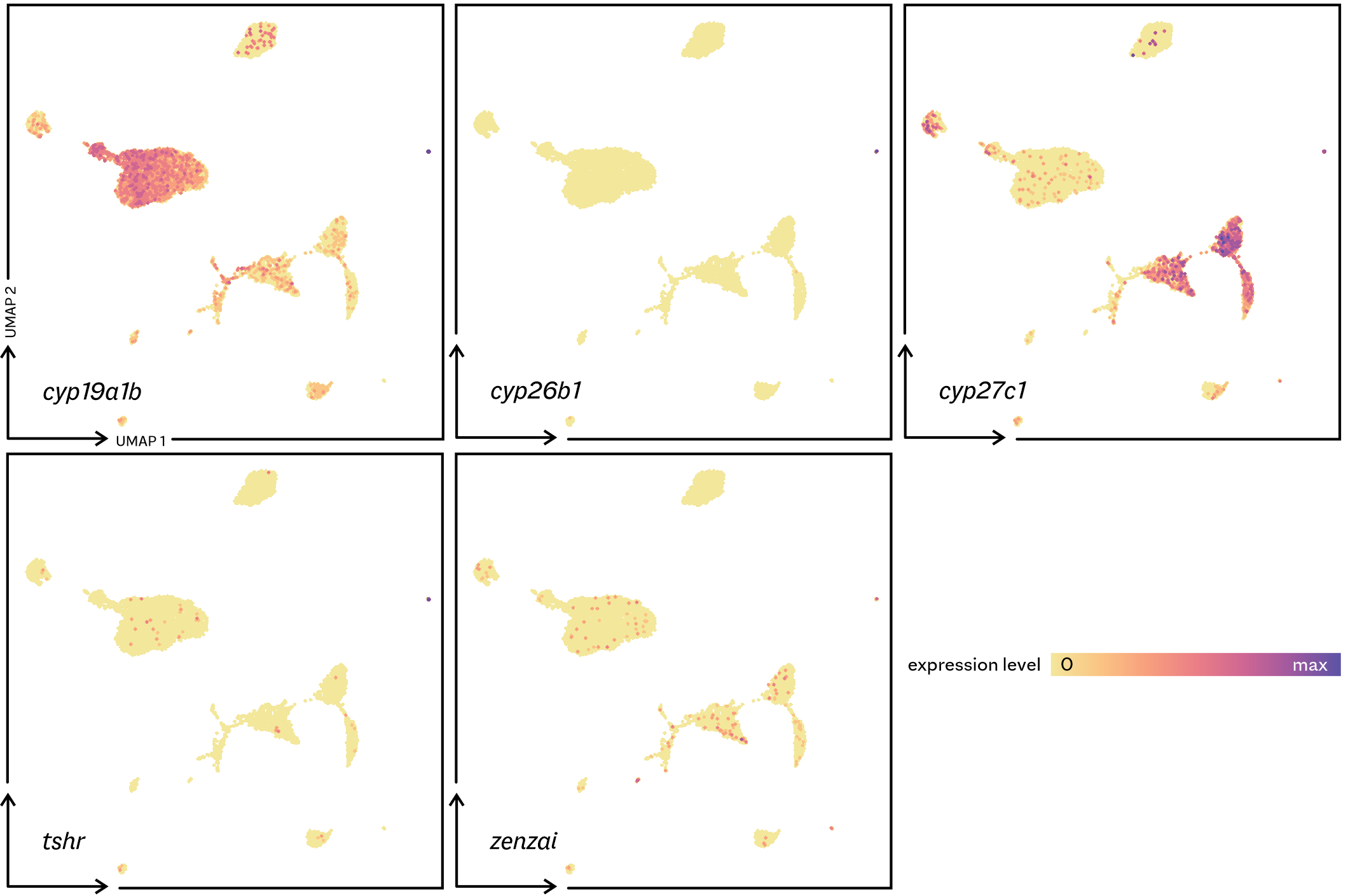

**Supplementary figure 12. Expression of selected cytochrome P450 and receptor genes in the scRNA-seq data.**

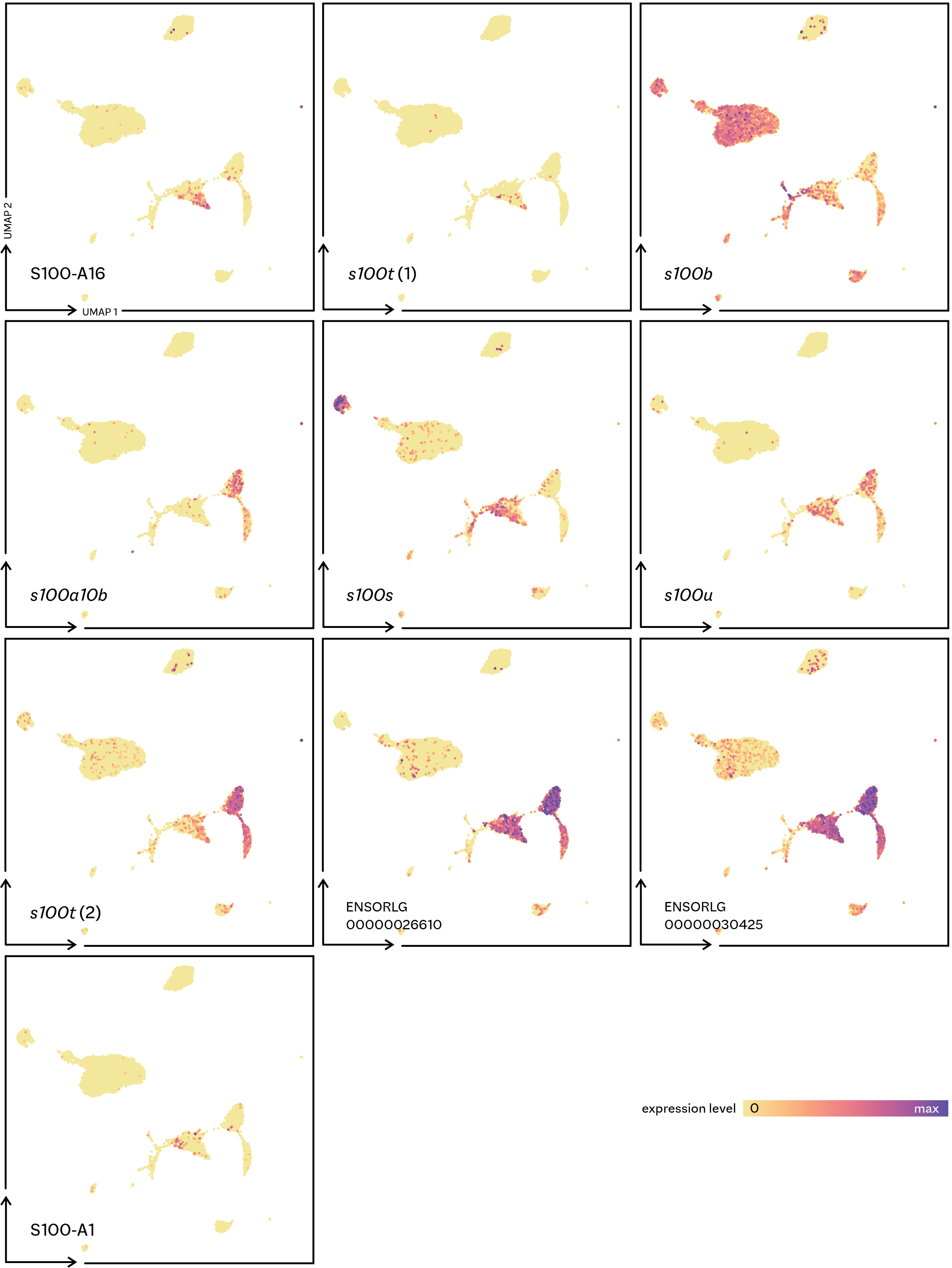

**Supplementary figure 13. Expression of *s100* paralogues often shows localization to subpopulations of the putative folliculo-stellate cell population.** Expressed medaka paralogues are: ENSORLG00000025488 (S100-A16), ENSORLG00000009724 (*s100t*), ENSORLG00000028907 (*s100b*), ENSORLG00000000531 (*s100a10b*), ENSORLG00000022495 (*s100s*), ENSORLG00000025692 (*s100u*), ENSORLG00000009724 (*s100t*), ENSORLG00000026610, ENSORLG00000030425, and ENSORLG00000023818 (S100-A1).

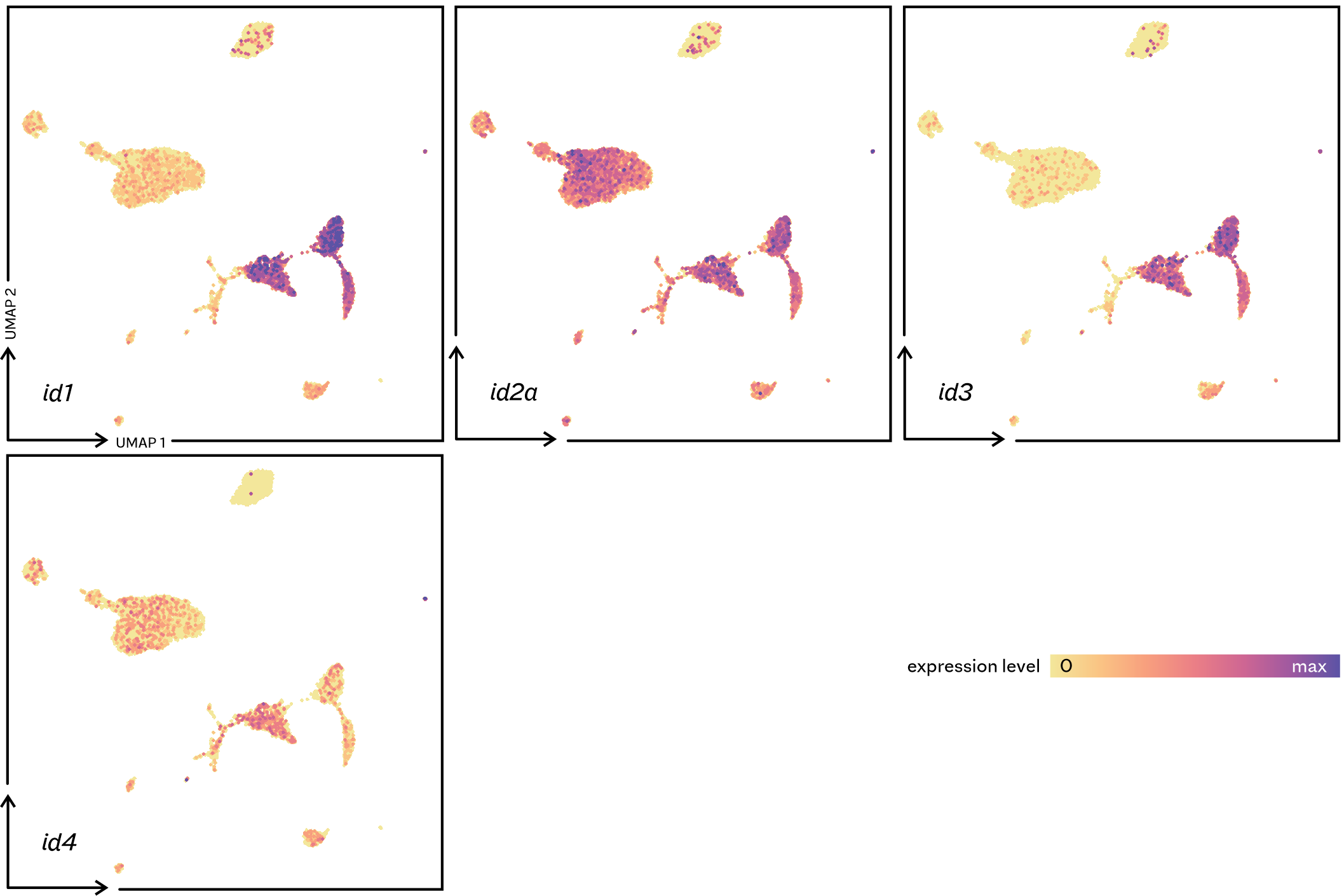

**Supplementary figure 14. Expression of *inhibitor of DNA binding* paralogues.**

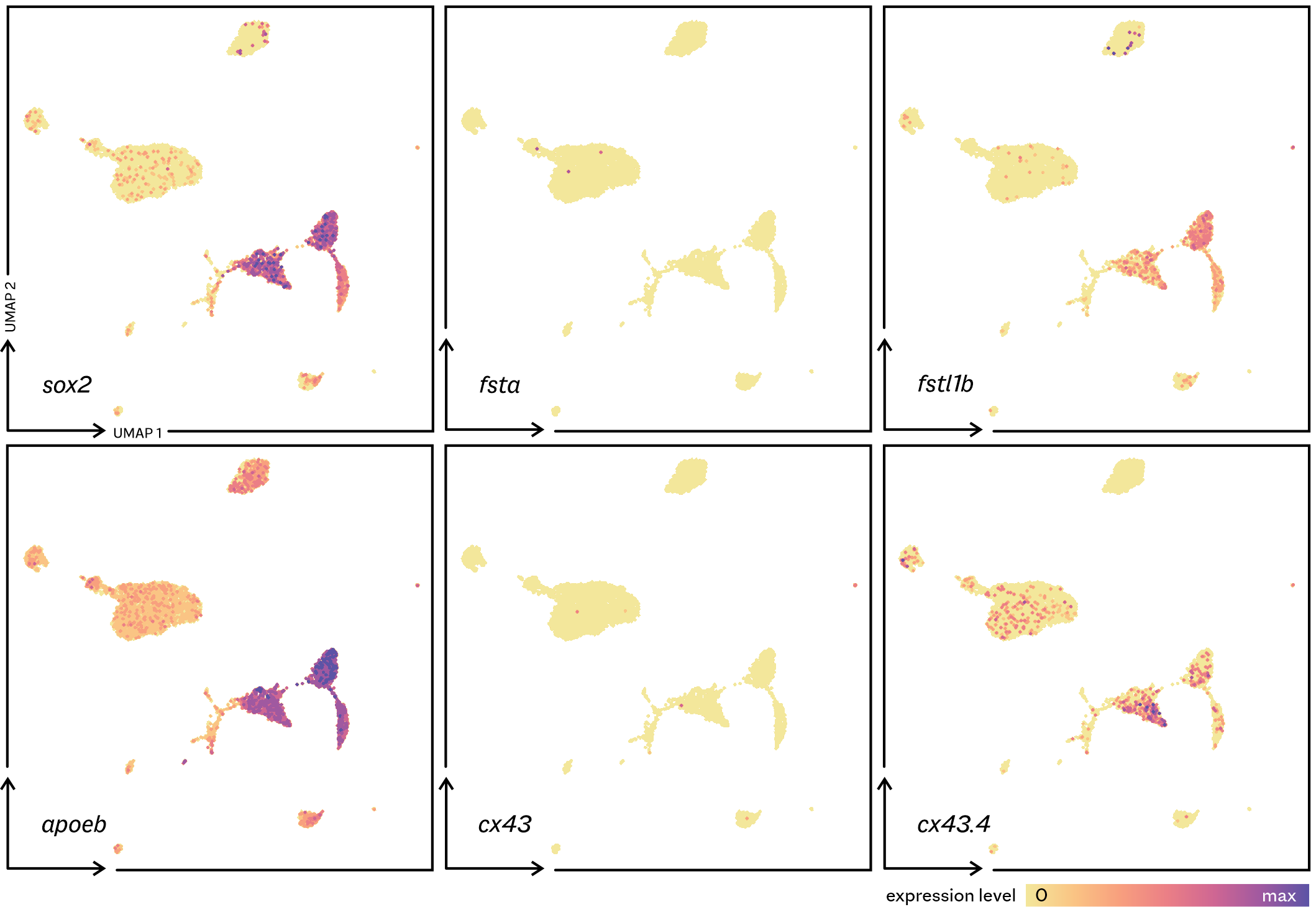

**Supplementary figure 15. Expression of follicolo-stellate and pituicyte marker genes.** *sox2* has two annotated paralogues in the medaka genome, with only ENSORLG00000001780 (shown here) expressed in the pituitary gland. Follistatin (*fsta*) is an established folliculo-stellate marker, however in medaka only follistatin-like (*fstl1b*) is expressed in the pituitary.

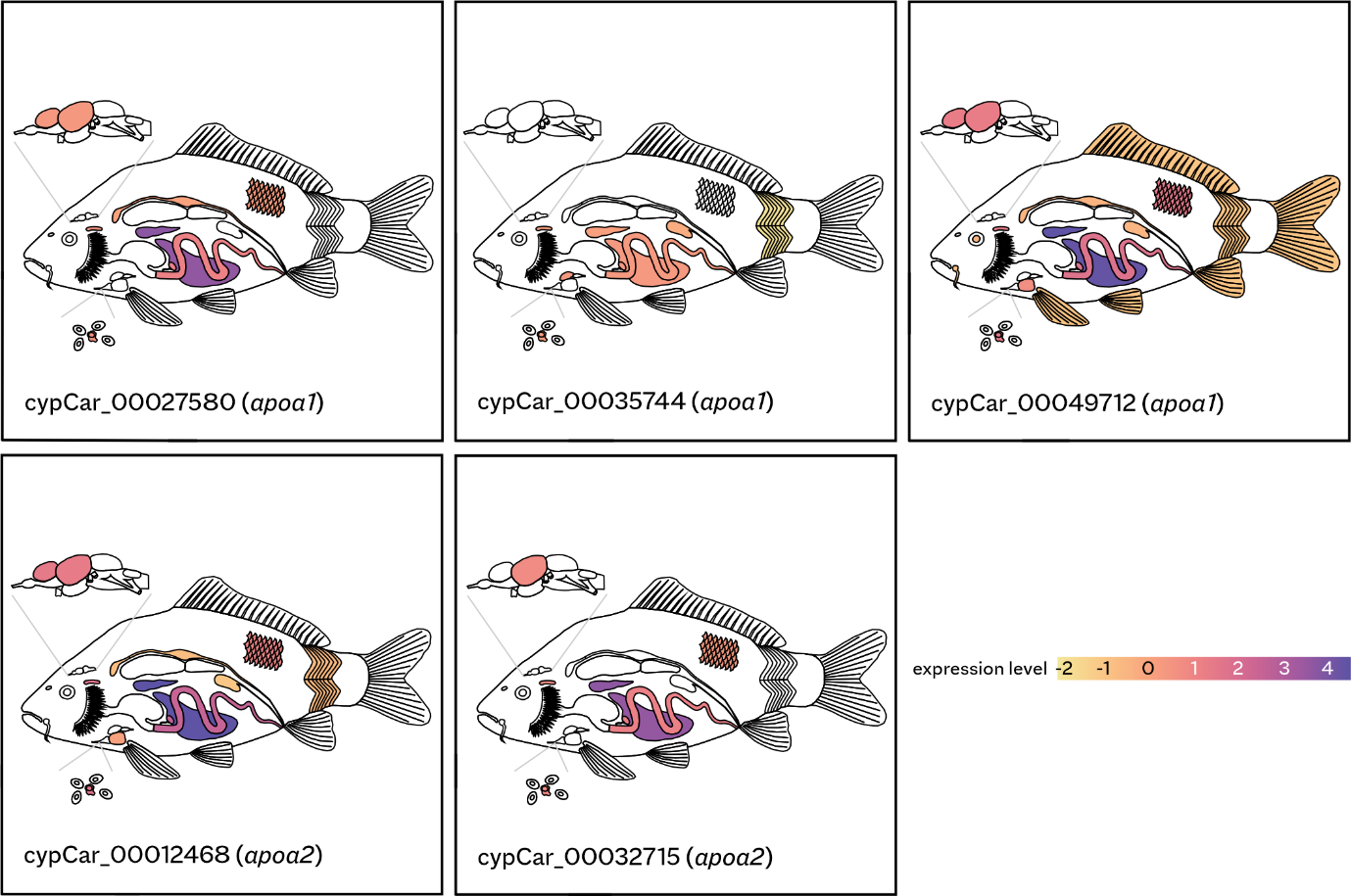
**Supplementary figure 16. Expression of *apoa1*/*2* orthologues in a carp (*Cyprinus carpio*) RNA-seq expression atlas.** Tissue expression (logarithmic scale) is shown for three *apoa1* paralogues (top) and two *apoa2* paralogues (bottom). The carp genome has experienced an additional whole-genome duplication compared to other teleosts, leading to more gene copies. High *apoa* expression is only found in the liver and the spleen, with low levels observed in many other tissues, including the brain (telencephalon and optic tectum). The pituitary gland does not display any *apoa* expression.

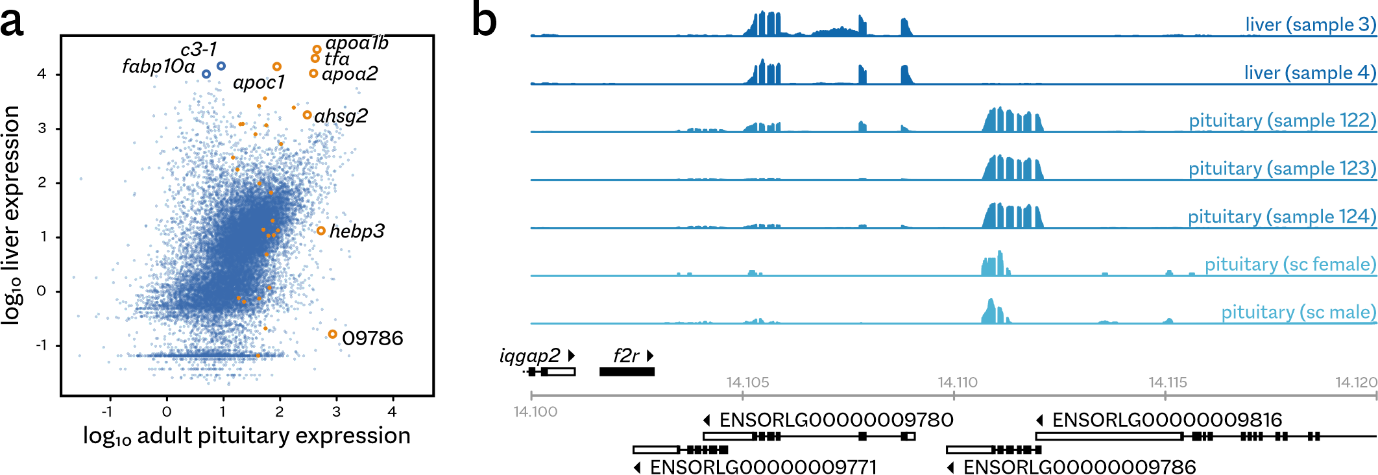

**Supplementary figure 17. Expression of liver-specific genes in the pituitary is not caused by liver tissue. a** Scatterplot showing the overall expression patterns in medaka pituitary and liver. Module 17 genes are highlighted in orange, with the most highly expressed annotated. Also annotated are the most highly expressed genes in the liver (open blue circles). The horizontal band of genes showing very low expression in liver is an artifact caused by the much higher sequencing depth for the liver samples. **b** Local gene expression for the orosomucoid locus on medaka chromosome 9. The locus consists of three unannotated local duplicates (ENSORLG00000009771/9780/9786), which are expressed at different levels in the pituitary and liver. A fourth gene, ENSORLG00000009816 (encoding single-stranded DNA binding protein 2) has a long 3’ UTR annotation that overlaps the first exon of orosomucoid ENSORLG00000009786. Expression profiles for each sample are scaled to the maximum local expression value. Note that different sequencing library preparation protocols affect expression patterns over a gene: for example, the 10x single-cell protocol (sc, lower two profiles) captures 3’ expression only.

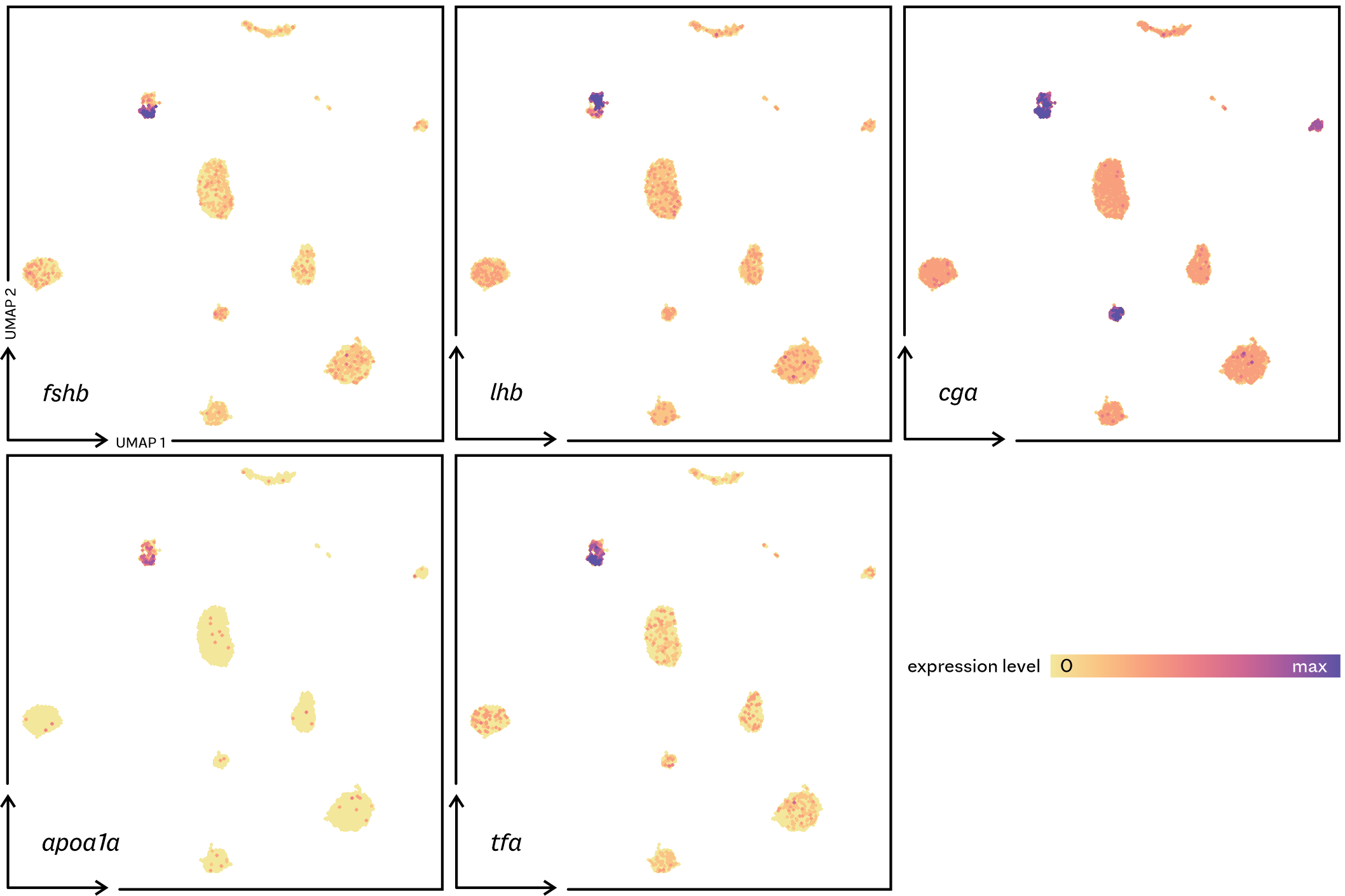

**Supplementary figure 18. In the zebrafish pituitary gland, *apoa1a* and *tfa* are expressed in gonadotrope cells (co-expressed with *fshb*, *lhb* and *cga*).**
